# Strong inhibition of insulin/IGF-1 signaling in early-mid adulthood compresses morbidity, but in later life accelerates aging

**DOI:** 10.64898/2026.04.15.718682

**Authors:** Bruce Zhang, Kuei Ching Hsiung, Reva Biju, Marybelle Cameron-Pack, Xiaoya Wei, Hannah Chapman, Melisa Kelaj, Aihan Zhang, Chiminh Nguyen Hong, Collin Y. Ewald, David Gems

**Author notes:** **Corresponding author:** David Gems, Institute of Healthy Ageing, and Department of Genetics, Evolution and Environment, University College London, Gower Street, London WC1E 6BT, United Kingdom.

## Abstract

Reduced insulin/IGF-1 signaling (IIS) can greatly extend lifespan in *C. elegans*. However, its effects on the duration of healthy life (healthspan) remain unclear, with several reports of either morbidity expansion or scaled effects, though none of morbidity compression. Moreover, life-extension by IIS reduction is particularly inter-individually variable within populations, confounding efforts to understand the intra-individual biology of such interventions. Here, we performed a longitudinal investigation at individual nematode resolution, of IIS reduction on aging-related health and lifespan, through temporally-controlled auxin-induced degradation (AID) of the DAF-2 insulin/IGF-1 receptor. Our results show how inter-individual variation in aging rate within control populations explains the complex demographic effects of age-specific DAF-2 AID on population lifespan. Strikingly, adult-limited IIS reduction causes an inter-individually homogeneous increase in lifespan (reducing Gompertz *α* rather than *β*) that is driven by healthspan expansion and compression of morbidity. Unexpectedly, cessation of DAF-2 AID in decrepit elderly individuals rejuvenates locomotory capacity and extends lifespan, showing that higher levels of IIS are optimal for health and survival towards the end of life. We also document a memory effect of transient IIS reduction during early adulthood, that is sufficient to fully extend lifespan (+189% median lifespan). Together, these findings demonstrate that both lifespan and healthspan can be maximized by appropriate temporal and directional modulation of IIS.

## Introduction

The insulin/IGF-1 signaling (IIS) pathway regulates key organismal processes including development, growth, metabolism and aging, across animal species including humans. In *Caenorhabditis elegans*, reduction of IIS can greatly extend lifespan, by as much as 10-fold (Friedman and Johnson, 1988, Kenyon et al., 1993, Ayyadevara et al., 2008), and weaker extensions have also been reported in other species (Vitale et al., 2019).

While IIS reduction robustly increases *C. elegans* lifespan, reports on the extent to which it increases the duration of healthy life (healthspan) are conflicting. For instance, some studies report increases in the duration of the later decrepit phase of life (gerospan), that increase the proportion of life in gerospan (expanded morbidity) (Huang et al., 2004, Bansal et al., 2015, Podshivalova et al., 2017, Churgin et al., 2017, Newell Stamper et al., 2018, Zhang and Gems, 2026), while others see scaled (i.e. proportionate) increases in healthspan and gerospan (Hahm et al., 2015, Statzer et al., 2022). Thus, either way, reducing IIS has not been found to increase healthspan more than gerospan (i.e. to compress morbidity). For any eventual translation of model organism biogerontology for human benefit, compression of morbidity would be desirable.

Less well studied is how IIS reduction increases lifespan variation within *C. elegans* populations (which are isogenic). IIS reduction typically produces larger extensions of the tail than of the shoulder of survival curves (Johnson, 1990, Samuelson et al., 2007, Zhang and Gems, 2026). This corresponds to a reduction in the rate parameter *β* of the Gompertz model of mortality rate (Gompertz, 1825), which reflects increased lifespan variation (Tuljapurkar and Edwards, 2011, Zhang and Gems, 2026). If and how this arises from increases in the inter-individual variation of aging remains unclear. This question exemplifies the wider challenge of understanding the relationship between aging at the individual and population level (Zhang et al., 2016, Zhao et al., 2017, Zhang and Gems, 2026, Zhang et al., 2026).

IIS reduction is typically achieved in *C. elegans* by mutation of genes within the pathway, commonly those encoding the DAF-2 insulin/IGF-1 receptor and its downstream AGE-1 phosphoinositide 3-kinase catalytic subunit, or by *daf-2* mRNA knockdown using RNA-mediated interference (RNAi). Another, more recent approach is the auxin-inducible degradation (AID) of DAF-2 protein, which enables rapid and stronger age-specific IIS reduction (Venz et al., 2021, Roy et al., 2022, Zhang et al., 2022, Molière et al., 2024). In brief, exposure to auxin (indole-3-acetic acid) of *C. elegans* of strains expressing degron-tagged DAF-2, such as RAF2181, induces proteasomal degradation of DAF-2 protein, thereby decreasing IIS. Importantly, by administering auxin from different ages, temporal effects of IIS on aging across the life course can be studied. For example, in this way it was shown that IIS reduction from advanced ages can still markedly extend lifespan (Venz et al., 2021), and potentially reverse some aging-related changes (Molière et al., 2024). Though informative, such studies do not take into account the large inter-individual variation in nematode aging, which can confound investigations of intra-individual processes (Zhang et al., 2016, Zhao et al., 2017, Zhang and Gems, 2026).

Here, we have employed the DAF-2 AID system to interrogate, within individually-tracked nematodes, the role of IIS in aging across the life course. Our longitudinal approach reveals an intrinsic inter-individual variability of aging rate within isogenic populations that can explain the variable effects of IIS reduction on individual lifespan. We show how this variation in aging rate impeded interpretability of earlier studies of anti-aging effects of IIS reduction in already-senescent animals (Venz et al., 2021, Molière et al., 2024). Surprisingly, accounting for such variation reveals that levels of IIS reduction that protect against aging in early adulthood instead *accelerate* aging during senescence, such that late-life cessation of DAF-2 AID extends lifespan and even modestly rejuvenates locomotory capacity. Importantly, DAF-2 AID extended healthspan markedly more than gerospan, demonstrating that IIS reduction, when optimally applied, can extend lifespan while compressing morbidity. Taken together, our investigation lays bare the complex roles of IIS in aging both across the life course and between isogenic population members.

## Results

### Age-dependent effects of IIS reduction on individual lifespan

We first assessed the effects on lifespan of initiating DAF-2 AID from different chronological ages. Auxin administration starting from later than L4 (final larval stage, designated as day 0 of adulthood) yielded progressively smaller increases in mean lifespan (Fig. 1a, S1a, Table S1), consistent with previous reports on the timing of *daf-2* action on lifespan (Dillin et al., 2002, Venz et al., 2021). These increases were no longer statistically significant (log-rank *p*>0.05) when auxin treatment was started from day 20 and later (Table S1). Extension of maximum lifespan was also reduced in magnitude with later auxin commencement (Fig. S2a). Nonetheless, these results recapitulated the earlier, remarkable observation (Venz et al., 2021) that auxin treatment even from very late ages, when many individuals have already died, can markedly extend lifespan (see e.g. D16 and D20 auxin, Fig. 1a). We also verified that auxin (in N2) and its solvent DMSO (in RAF2181) do not themselves affect lifespan (Fig. S2c–d), as previously shown (Venz et al., 2021).

**Fig. 1.**
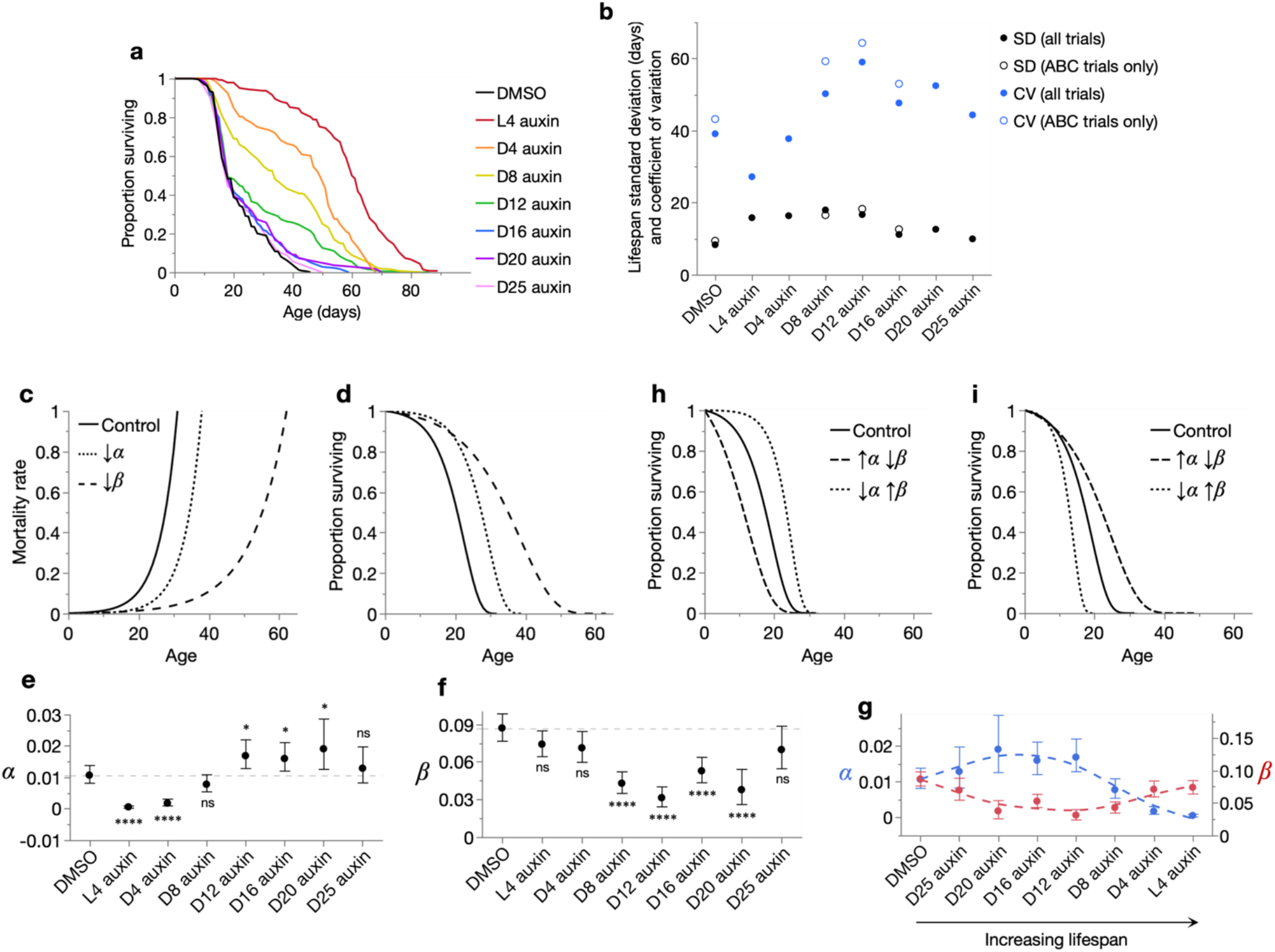
Effects of age-specific DAF-2 AID on lifespan and gerodemography. (**a**) Kaplan-Meier survival curves for the age-specific DAF-2 AID and DMSO-only control cohorts (pool of all trials; see Table S1 for lifespan statistics). (**b**) Standard deviation (SD) and coefficient of variation (CV = SD/mean, a measure of relative rather than absolute variability) of lifespan of the control and DAF-2 AID cohorts, for either the pool of all trials or only trials in which locomotory capacity was scored (see Table S1). (**c**– **d**) Effects of decreasing the Gompertz parameters *α* and *β* on mortality rate (**c**) and corresponding survival proportion (**d**) (formal scheme). Control: *α, β*: 0.002, 0.2; ↓*α*: *α, β*: 0.0005, 0.2; ↓*β*: *α, β*: 0.002, 0.1. The Gompertz equation is: *μ(x)=αe*^*βx*^, where *μ(x)*=mortality rate, *x*=age, *e*=Euler’s number, *α*=scale parameter, and *β*=rate parameter. (**c**) *α* and *β* respectively control the scale and rate of exponential mortality rate increase. (**d**) *α* and *β* approximately shift and stretch, respectively, the survival curve along the x-axis. That is, inter-individual lifespan variation is increased more by reductions in *β* than *α* (Tuljapurkar and Edwards, 2011). (**e**–**f**) Effect of age-specific DAF-2 AID commencement on the Gompertz parameters *α* (**e**) and *β* (**f**), for the pool of all trials. Statistical significance of parameter differences (via likelihood ratio tests) between cohorts (versus DMSO) is indicated, displaying 95% confidence intervals. (**g**) Gompertz parameters from the control and DAF-2 AID cohort data arranged in order of increasing lifespan, which reveals an inverse, Strehler-Mildvan correlation between the parameters. (**h**–**i**) Two forms of survival curve transformation characteristic of the Strehler-Mildvan correlation: (**h**) rectangularization and de-rectangularization, and (**i**) triangularisation and de-triangularisation (formal scheme) (Zhang et al., 2026). The parameters are: (**h**) Control: *α*=0.002, *β*=0.2; **↑*α*↓*β***: *α*=0.012, *β*=0.14; **↓*α*↑*β***: *α*=0.0001, *β*=0.29, and (**i**) Control: *α*=0.002, *β*=0.2; **↑*α*↓*β***: *α*=0.003, *β*=0.1; **↓*α*↑*β***: *α*=0.0012, *β*=0.37.

Such age-specific responses to DAF-2 AID could reflect age-specific roles of IIS, or population heterogeneity that changes with age or, most plausibly, a combination of the two. Individual nematodes age at different rates (hence their different lifespans) (Zhang et al., 2016), such that as populations age, individuals within the population exhibit increasingly different biological ages at the same chronological age. Consequently, one might expect different responses to IIS reduction when started from the same chronological age, particularly in later life.

Consistent with this, median lifespan was increased only by auxin administration initiated prior to day 10 (Fig. S2b) despite increases in mean and maximum lifespan given later treatments (Fig. S2a, Table S1). Moreover, excluding pre-auxin treatment deaths altered the effect of later auxin treatments, increasing the magnitude of mean lifespan extension by DAF-2 AID from day 20 (from +14%, *p*=0.055, to +22.3%, *p*=0.003), but not (even decreasing it) for DAF-2 AID from days 16 and 25 (from +7.9% to +4.3%, and +3.5% to +3%, respectively) (Tables S1–2). These observations show that population members respond differently to treatment, suggesting that the distribution of inter-individual variation in biological age at the time of DAF-2 AID commencement determines its effects on population survival.

Importantly, unless specified otherwise, we include pre- and near-intervention deaths in all of our demographic analyses as these individuals belong to a continuous, inter-individual spectrum of intervention efficacy, ranging from maximally effective to wholly ineffective. How non- and weak-responding individuals impact population-level measures is complex, especially when their presence and prevalence are unknown (as in most interventions).

### Adult-limited reduction of IIS reduces Gompertz *α* but not *β*

Inter-individual variation in lifespan responses is captured in the shapes of survival curves. DAF-2 AID from L4 approximately shifted the survival curve rightward, whereas progressively later DAF-2 AID increasingly diminished the lifespan gains in the curve shoulder, as expected (Fig. 1a). Instead, later DAF-2 AID lengthened the survival curve tail, thereby increasing lifespan variation (standard deviation and coefficient of variation; Fig. 1b). This increased lifespan variation was not solely due to the presence of pre-DAF-2 AID deaths, as it persisted when such deaths were excluded (Fig. S1b). Therefore, commencing DAF-2 AID after early adulthood produces not only weaker but more heterogeneous effects on the lifespans of individuals.

We next studied the lifespan variation of our cohorts using the Gompertz model of mortality rate (*μ(x)=αe*^*βx*^) (Gompertz, 1825) (Fig. 1c) and its corresponding survival function (Fig. 1d). Here the scale parameter *α* controls the horizontal position of the survival curve (i.e. when mortality noticeably begins), while the rate parameter *β* controls its steepness and captures the amount of lifespan variation (smaller *β* = reduced survival curve steepness and greater lifespan variation) (Tuljapurkar and Edwards, 2011); note that the earlier interpretation of *β* as a metric of biological aging rate is largely erroneous, at least in *C. elegans* (Zhang and Gems, 2026). The Gompertz model provided an appropriate fit to each cohort (Fig. S1c, Table S3: parametric Anderson-Darling goodness-of-fit tests). While more complex models may offer better fits (Vanfleteren et al., 1998, Johnson et al., 2001, Mulla et al., 2023), especially in the mid-late DAF-2 AID cohorts, Fig. S1c, the Gompertz model is more widely used across conditions and species, mathematically parsimonious (with only 2 parameters), and effectively captures changes in lifespan variation in our cohorts (Fig. S1d).

Interestingly, auxin treatment from L4 strongly decreased *α* but not *β* (Fig. 1e–f, Table S1), corresponding to the largely parallel right-shift of the survival curve (Fig. 1a); this contrasts with many earlier findings that IIS reduction decreases *β*, as in the landmark *age-1(hx546)* mutant study (Johnson, 1990). In later DAF-2 AID, decreases in *β* emerged and grew progressively more marked (Fig. 1f), consistent with increased lifespan variation resulting from variable DAF-2 AID efficacy across population members. However, the magnitude of *β* reduction decreased with very late DAF-2 AID (∼day 16 onwards) (Fig. 1f), in line with diminishing extensions of the survival curve tail (Fig. 1a). Meanwhile, reductions in *α* also diminished with later DAF-2 AID (Fig. 1e), in line with diminishing extensions of the survival curve shoulder (Fig. 1a). This magnitude reduction continued in treatments later than day 8, even changing into modest increases in *α* relative to DMSO control values (Fig. 1e).

### Late onset IIS reduction generates a spectrum of Strehler-Mildvan correlations

Notably, the Gompertz parameters changed inversely as lifespan increased across the cohorts (Fig. 1g). The smallest lifespan extensions (by very late DAF-2 AID) increased *α* while decreasing *β*, while further lifespan extensions (by late to progressively earlier DAF-2 AID) reversed this pattern: decreasing *α* while increasing *β*. This reveals a Strehler-Mildvan (S-M) correlation in the age-specific effects of IIS reduction on mortality rate, i.e. an inverse relationship between *α* and *β*, which is often observed across human and animal populations (e.g. Strehler and Mildvan, 1960, Shen et al., 2017). Importantly, the appropriate fits of the Gompertz model to our data (Fig. S1c–d, Table S3) indicate that this particular S-M correlation is not a fitting artifact (Tarkhov et al., 2017) but reflects biologically reproducible changes in lifespan.

S-M correlations reflect the rectangularization/de-rectangularization (Fig. 1h) or triangularization/de-triangularization of survival curves (Fig. 1i) (Zhang et al., 2026). Here both types of S-M correlation were observed. Very late DAF-2 AID triangularized the survival curve (extending its tail) by increasing *α* and decreasing *β* (Fig. 1a, g). Progressively earlier DAF-2 AID increased the magnitude of triangularization, and which gradually transitioned into rectangularization (extending the survival curve shoulder). We and others have reported that life-extending S-M correlations (rectangularization and triangularization) reflect inter-individually variable responses to interventions (Kirill et al., 2024, Simon et al., 2026, Zhang et al., 2026). This supports the view that age-specific DAF-2 AID (relative to the DMSO control or other DAF-2 AID timepoints) is an inter-individually variable intervention. In particular, DAF-2 AID from after day 8 of adulthood triangularized survival relative to the DMSO control (Fig. 1e– f), indicating increases in the within-population heterogeneity of aging.

### Emergence of individual variation in locomotory capacity during aging

To confirm whether and explore how inter-individual variation in biological age causes such age-specific DAF-2 AID effects on lifespan, we examined the aging-related health of all population members at each age of DAF-2 AID commencement. Locomotory capacity was scored every 2–3 days as described (Zhang and Gems, 2026), adapted from an established classification system (Hosono et al., 1980, Herndon et al., 2002). In brief, nematodes were assigned to movement classes based on their response to a physical stimulus: A class (sinusoidal locomotion), B class (non-sinusoidal locomotion), and C class (moving but non-locomotory). The days spent in each class we will refer to as A-span, B-span, and C-span, which sum to lifespan.

As expected, in early adulthood (before day 8), the DMSO control population was highly homogeneous with all individuals in the A (healthiest) class (Fig. 2a–b). From day 8 onwards, the population became increasingly heterogeneous as individuals began to enter B and C classes (and to die). These changes appeared chronologically earlier in shorter-lived population members, in agreement with their presumably faster and/or earlier aging.

**Fig. 2.**
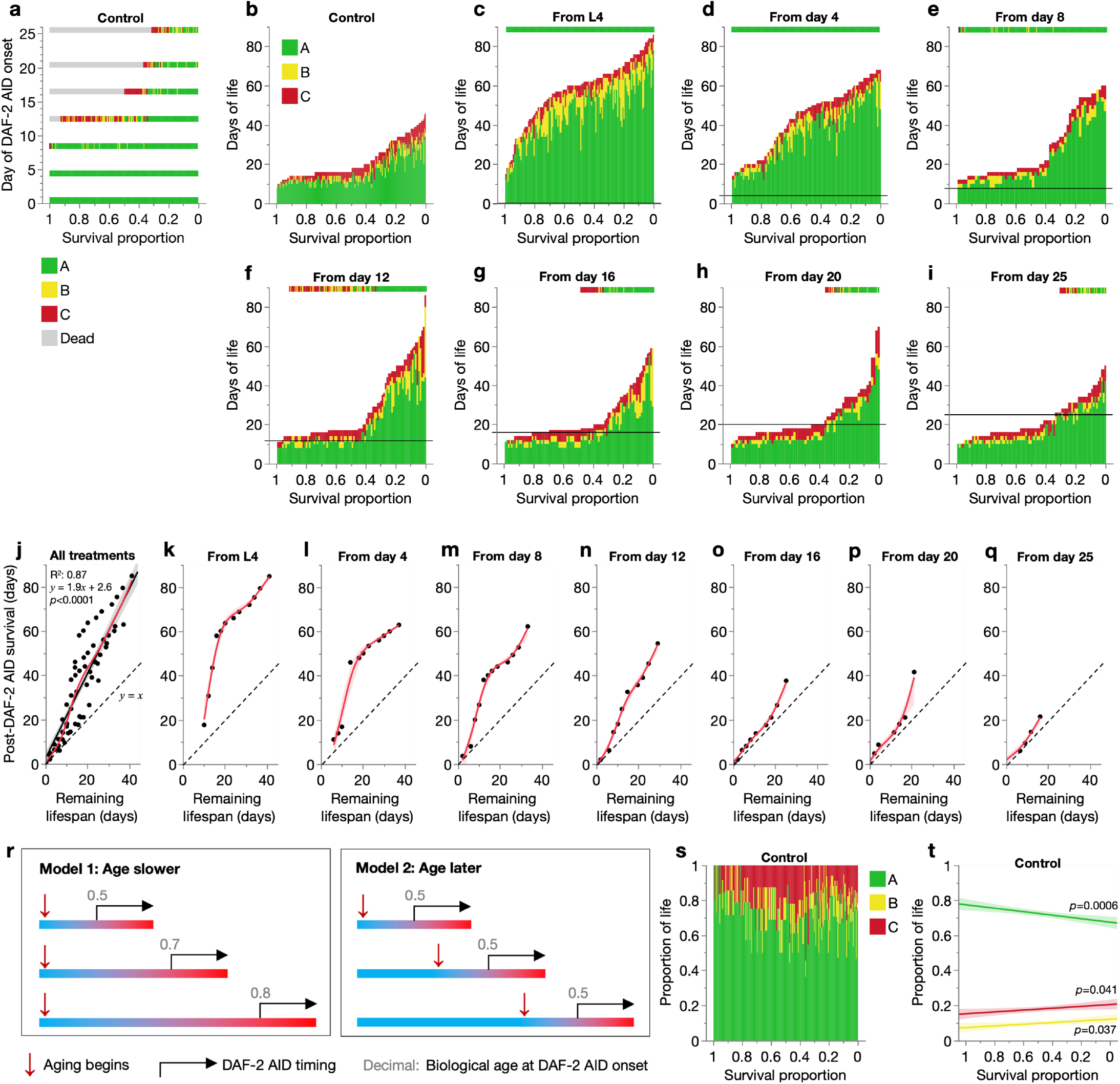
Effects of inter-individual variation in aging rate on responses to DAF-2 AID. (**a**–**i**) Vertically stacked days of life spent by each individual in A, B and C locomotory classes, ordered along the x-axis by decreasing survival proportion (increasing lifespan). Each vertical bar represents one individual. (**a**) Cross-sectional slices of the DMSO control event history chart (**b**), corresponding to the days that auxin treatment was commenced. (**c**–**i**) Auxin-treated cohorts; horizontal black lines indicate the day of auxin commencement. A cross-sectional slice of the DMSO control event history chart corresponding to the day of auxin commencement is displayed above each auxin-treated cohort for comparison. Note that the heterogeneity in movement class at the age of DAF-2 AID commencement increases from **c** to **i**. (**j**– **q**) Effect of remaining DMSO control lifespan at the age of auxin commencement on days survived after DAF-2 AID commencement in the auxin-treated cohorts, for all auxin-treated cohorts combined (**j**) or separate (**k**–**q**). To plot this relationship, remaining lifespan of control animals and post-DAF-2 AID survival of treatment animals were related to one another by survival proportion, which was grouped into 0.05-unit bins (i.e. 20 survival proportion bins between 1 and 0). Remaining control lifespan and post-DAF-2 AID survival of individuals falling within these bins were averaged to yield one value per bin, and these were plotted as above (depicted datapoints) and their relationship summarised with a non-linear smoother (spline method with lambda of 0.05, showing 95% confidence regions). The dashed lines represent a 1:1 relationship between the axis variables, which would be expected if DAF-2 AID has no effect on lifespan, and the red curves are spline smoothers (lambda=0.05). All data in this figure are from the pool of Trials 1–5, with censored individuals excluded. (**r**) Two potential explanations of lifespan variation within populations. Model 1 (left), where longer-lived population members age at a slower rate throughout life, and Model 2 (right), where they age at the same rate but starting from a later age. Each bar represents the life course of an individual; blue: healthspan, red: gerospan, bar length: lifespan. (**s**– **t**) Relationship between survival proportion and the proportion of life spent in A, B and C locomotory classes for DMSO-treated control individuals, displayed as a life-history chart in **s** and linear regression in **t**. Regression F-test *p*-values are labelled in **t** for each locomotory class, displaying 95% confidence regions.

Notably, this age-increase in inter-individual variation in locomotory health (i.e. biological age) agrees with the age-specific effects of DAF-2 AID on survival curve shape (and Gompertz parameters), as follows. IIS reduction in a young (L4), pre-senescent (more homogeneous) population (Fig. 2a, c–d) produced similar longevity responses across individuals, shifting the survival curve rightward and decreasing *α* only (Fig. 1a, e–f). Initiation of IIS reduction in older (increasingly heterogeneous) populations (Fig. 2a, e–i) produced increasingly variable longevity responses across individuals, thus only extending the survival curve tail and decreasing *β* instead (Fig. 1a, e–f).

### Variation in aging rate determines individual lifespan responses to IIS reduction

Next, we characterized in greater detail the emergence of inter-individual differences in biological age, and their effects on responses to IIS reduction. A simple hypothesis is that longer-lived individuals age more slowly (and/or later), and are thus biologically younger than their shorter-lived counterparts at any given chronological age in adulthood. If the ability of IIS reduction to ameliorate aging depends on biological age, this could explain the inter-individually variable effects of later onset DAF-2 AID on lifespan.

Consistent with this hypothesis, even during the early adulthood period (L4–day 8) when all population members are in the healthiest A class, later DAF-2 AID produced smaller lifespan increases (Fig. 1a, 2a–e). This suggests that despite exhibiting visibly youthful locomotion, A class animals have already begun to age, here detectable in their declining responsiveness to IIS reduction. In line with this, DAF-2 AID initiated later in this time window already began to increase lifespan variation (Fig. 1b) and decrease *β* (Fig. 1f), indicating that biological ages begin to vary (due to different individual aging rates) well before visible locomotory defects appear. This is consistent with known senescent changes during the first few days of adulthood in *C. elegans* (Huang et al., 2004, Labbadia and Morimoto, 2015, E et al., 2018).

To further investigate how biological age at treatment onset determines the strength of DAF-2 AID effects, we examined the relationship between control and DAF-2 AID animals of survival following treatment commencement, across all DAF-2 AID cohorts (Fig. 2j). Post-treatment survival increased linearly with remaining control lifespan, with a slope greater than 1 (Fig. 2j), showing that *within* these populations, reducing IIS at younger biological ages increases lifespan more. Post-treatment survival increased with a slope of 1.9; thus, on average across the cohorts, DAF-2 AID approximately doubles the remaining lifespan of any treated individual.

Considering these relationships for each age-specific DAF-2 AID treatment separately (Fig. 2k–q), one also observes gradients steeper than 1 (Fig. 2e). This includes treatments starting before ∼day 8 (notably, even as early as L4), supporting the view that responsivity to IIS reduction declines, and at different rates between individuals, right from the start of adulthood. Together, these findings imply that during A-span, progressive changes occur (pre-pathological and/or pathological) that precede and give rise to healthspan decline and death, and which are promoted by IIS. Hence, the later in A-span DAF-2 AID is initiated, the smaller the increase in lifespan. This suggests that aging rate is slower throughout adulthood in longer-lived individuals.

Notably, with earlier treatment initiation (L4–day 8), the relationship is bimodal, with an initially greater slope that lowered when remaining lifespan reached ∼day 20 (Fig. 2k-m). The steeper first stage may correspond to the efficacy of reducing IIS in individuals that otherwise die prematurely due to pharyngeal infection, which typically die within this window (days 10–20) (Zhao et al., 2017). The second flatter stage, whose slope of ∼1 indicates no benefit of earlier treatment, may reflect already maximal treatment efficacy in longer-lived individuals that die without pharyngeal infection. That is, biological age at treatment onset is sufficiently low in these longer-lived individuals, such that earlier treatment offers no further benefit.

Comparing the age-specific DAF-2 AID cohort plots (Fig. 2k–q), a major difference between them is evident. For a given remaining control lifespan at DAF-2 AID onset (x-axis), the corresponding post-DAF-2 AID survival (y-axis) differs greatly. For instance, for a remaining lifespan of 20 days, post-DAF-2 AID survival ranged from ∼60 days (treatment from L4; Fig. 2k) to only ∼20–25 days (treatment from day 16 or later; Fig. 2o–q). This means that not all individuals with the same remaining lifespan respond equally to DAF-2 AID. This finding allows us to distinguish between two explanatory models of lifespan variation within populations, where longer-lived population members experience aging either predominantly more slowly or later, compared to their shorter-lived counterparts (Fig. 2r). Our finding that lifespan responses to DAF-2 AID differ despite the same post-treatment lifespan implies that biological age differs at treatment onset amongst such individuals. This further supports the view that aging is a continuous process throughout adulthood, that is slower (Fig. 2r, left) rather than delayed (Fig. 2r, right) in longer-lived population members.

Consistent with this model, the proportion of life spent in each locomotory class was relatively constant across control population members (Fig. 2s–t), suggesting an approximate stretching out of the overall aging process in longer-lived members. The proportion of life in A-span was even moderately reduced in longer-lived members (Fig. 2t), further arguing against a delaying of aging, which would be expected to increase it (Fig. 2r, right).

In short, these results imply that lifespan variation within the control population arises from inter-individual variation in aging rate. Aging (here measured as declines in locomotory capacity and efficacy of DAF-2 AID) begins in early adulthood, proceeds more slowly throughout life in longer-lived individuals, and is accelerated by IIS throughout adulthood. Critically, this variation in individual aging rate produces variation in biological age, which increases with chronological age; this explains the inter-individually variable lifespan responses to IIS reduction that depend on the age of treatment onset. Specifically, the life-extending effects of DAF-2 AID are largely exerted in A stage animals, but effects grow progressively weaker as A-span is traversed.

### IIS reduction by DAF-2 AID extends healthspan and compresses morbidity

Is gerospan, the period of aging-related morbidity, extended in long-lived *C. elegans* with reduced IIS? This question has been the subject of some controversy. Expressed as a proportion of overall lifespan, both expansion and proportional scaling of gerospan have been reported (Bansal et al., 2015, Hahm et al., 2015, Podshivalova et al., 2017, Statzer et al., 2022, Zhang and Gems, 2026). These studies mostly use reduction-of-function (rf) *daf-2* mutations that reduce IIS throughout life (including development). To investigate this further, we assessed how IIS reduction by DAF-2 AID affects healthspan and gerospan, which has not yet been investigated longitudinally.

Surprisingly, DAF-2 AID extended locomotory healthspan (A-span) at all ages of treatment initiation, with little or no increase in locomotory gerospan (B-span + C-span) (Fig. 3a–g). This resulted, in the main, in a compression of morbidity as a proportion of total adult lifespan (Fig. 3h–n), in contrast to earlier, *daf-2(rf)* mutant studies. Consistent with the inter-individual variation in aging rate characterised earlier (Fig. 2), these changes were increasingly restricted to longer-lived population members (lower survival proportions), and reduced in magnitude, when DAF-2 AID was initiated at later chronological ages. This compression (rather than proportional scaling or expansion) of morbidity likely reflects the stronger and/or adult-restricted IIS reduction, achieved here by using the DAF-2 AID system. The greater strength of IIS reduction by DAF-2 AID is indicated by the developmental arrest (as dauers) of all larvae undergoing DAF-2 AID during development, even at developmentally-permissive lower temperatures, in contrast to *daf-2* mutation and RNAi (Venz et al., 2021). Moreover, given the positive correlation between the severity of the constitutive dauer formation (Daf-c) phenotype and lifespan extension amongst *daf-2* alleles (Gems et al., 1998), the greater longevity resulting from early DAF-2 AID than the strong *daf-2(e1370)* allele (Table S1) (Venz et al., 2021, Zhang et al., 2022) is also in line with greater IIS reduction in the former.

**Fig. 3.**
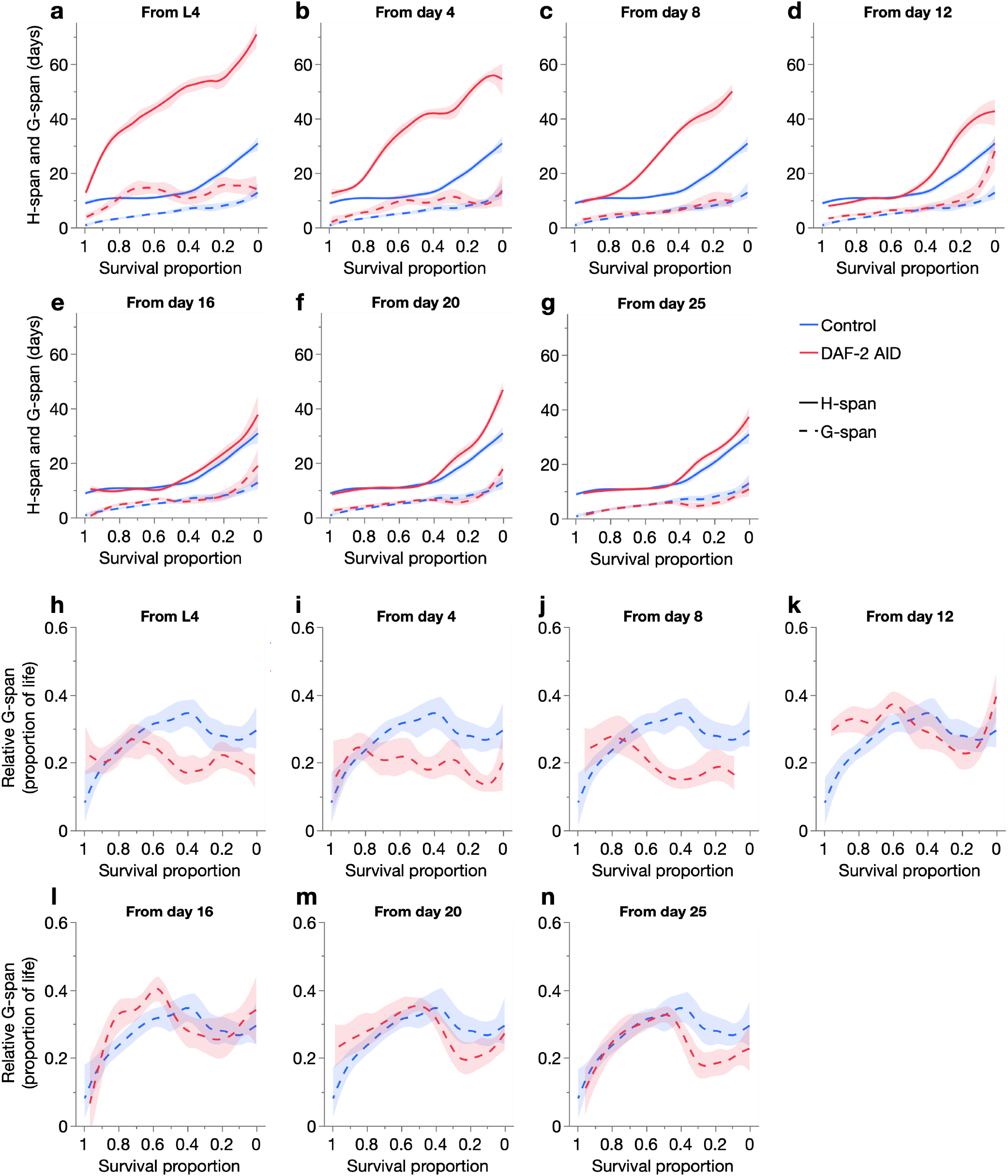
Effects of age-specific DAF-2 AID on locomotory healthspan and gerospan. Effect of age-specific DAF-2 AID on (**a***–***g**) absolute (days in) healthspan (H-span) and gerospan (G-span) and (**h**–**n**) relative (proportion of life in) G-span, compared to the DMSO-treated control, across individuals of each population as ordered by survival proportion (left: shorter-lived; right: longer-lived).

Despite their extended healthspan, we noticed that DAF-2 AID animals sometimes exhibited little immediate response to physical stimulus (e.g. gentle touch with a platinum wire), but after a short delay (5–10 sec) displayed normal A-class locomotion (and were therefore scored as such). Other forms of reduced motility have been previously noted with DAF-2 AID (Roy et al., 2022) and in class 2 (more pleiotropic) *daf-2(rf)* mutants (Gems et al., 1998, Hahm et al., 2015), and interpreted as a recapitulation of dauer immotility. However, a possibility in the present case is that such atypical A-class animals, which we will refer to as A’ animals (exhibiting A’ locomotion), are in a later and subtly senescent stage of A-span. The remaining portion of A-span, with un-delayed responsivity to physical stimuli, we will refer to as the canonical A-span (i.e. A-span = canonical A-span + A’-span).

To investigate whether A’ locomotion represents a true senescent state, we further characterized it in animals undergoing DAF-2 AID from L4. As expected, a higher proportion of these animals exhibited A’ locomotion at some point in their life (49/66: 74%) than untreated controls (10/71: 14%) (chi-squared *p*=0.0005) (Fig. 4a– b). Notably, A’ locomotion appeared only in later adulthood (after ∼day 15 in DAF-2 AID animals; Fig. 4b) and increased in incidence with age, but largely preceding the transition from H-span (A class) to G-span (B or C class). This suggests that A’ locomotion represents an intermediary stage of aging between the canonical A and B class states, unlike the behavioral effects of IIS reduction that can already be observed in early adulthood (Gems et al., 1998, Roy et al., 2022). Consistent with this, A’ animals did not revert at a higher rate to canonical A class when IIS was restored by stopping DAF-2 AID (Fig. 4c). Importantly however, DAF-2 AID animals frequently switched back and forth between A’ and canonical A locomotion (Fig. 4b), likely reflecting an extended period of gradual senescence, akin to the mid-life plateau in severity of several senescent pathologies (Ezcurra et al., 2018, Kern et al., 2025). Given this, we define A’-span and canonical A-span as the sum of total days observed in these locomotory classes, rather than continuous, uninterrupted stages of life.

**Fig. 4.**
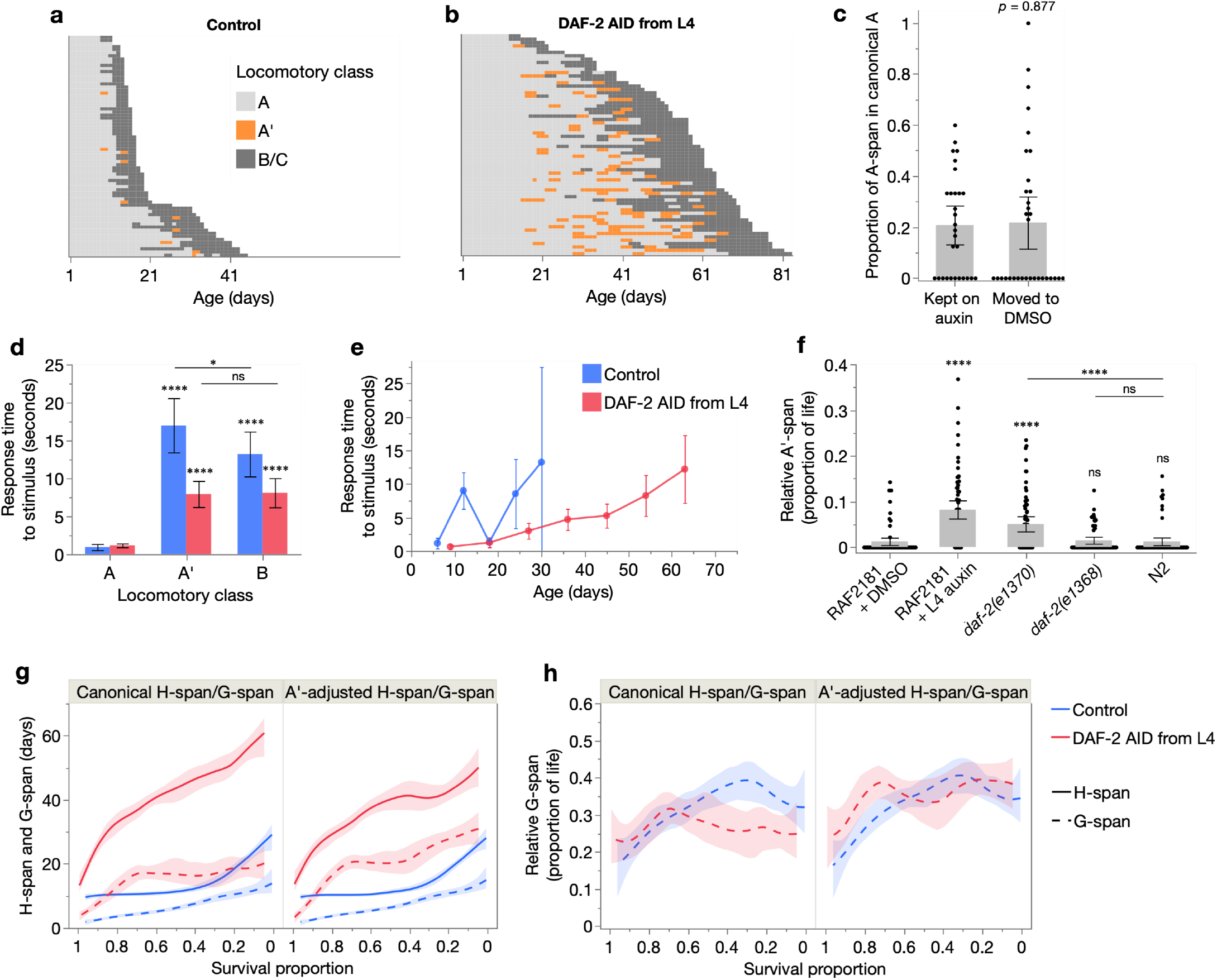
Characterisation of A’ locomotion. (**a**–**b**) Event history charts of the locomotory class of each individual throughout life, for those treated from L4 with DMSO only (**a**) or auxin (**b**). Each row represents one individual. (**c**) Mean proportion of post-transfer day A-span that was in only canonical A-span (rather than A’-span), for animals kept on auxin or taken off it (moved to DMSO alone) (compared with a two-tailed Student’s t-test). Animals were treated with auxin from day 12 and transfer day was when A’ locomotion was first observed (animals were scored every 1-2 days). (**d**–**f**) Mean time taken to begin locomoting away from one gentle touch to the tail with a platinum wire, for (**d**) A, A’, and B class animals, averaged from measurements taken throughout life, and (**e**) with age, for animals treated from L4 with DMSO only or auxin. (**f**) Mean relative A’-span (proportion of life spent in the A’ state) of different cohorts, from the pool of two trials. In **c**–**f**, 95% confidence intervals are shown, and statistical comparisons (two-tailed Student’s t-tests) are made with the first condition where of interest, except where indicated otherwise; ns *p* > 0.05, * *p* ≤ 0.05, ** *p* ≤ 0.01, *** *p* ≤ 0.001, **** *p* ≤ 0.0001. (**g**–**h**) Effect of auxin from L4 on (**g**) absolute H-span and G-span and (**h**) relative G-span, using either the canonical definitions (H-span = A-span, G-span = B+C-span) or A’-adjusted definitions (H-span = canonical A-span, G-span = A’+B+C-span). These relationships were fit with a smoother (spline method, lambda=0.05) displaying 95% confidence regions. All panels except **c** are from the pool of Trials 3 and 5, in which A’ was scored (sample sizes in Table S1).

Mean response time (to locomote away from a physical stimulus) of A’ animals was considerably longer than that of canonical A animals, and comparable to that of senescent B class animals (Fig. 4d). Consistent with these results, mean response time (which increases during aging) of all population members was lower throughout life in the DAF-2 AID than control cohort, and took twice as many days to reach the same maximum (Fig. 4e).

Mean relative (proportion of life in) A’-span was also higher in DAF-2 AID animals (Fig. 4f), indicating a disproportionate expansion of this senescent life stage. Notably, relative A’-span was also increased in *daf-2(e1370)*, but not *daf-2(e1368)* and N2, consistent with dauer-like motility reductions in the former only (Gems et al., 1998). However, our findings suggest that the reduced motility of *daf-2(e1370)* at 20°C may reflect true senescence, in the form of an expanded A’-span. Consistent with this, A’ locomotion in *daf-2(e1370)* was confined to later adulthood, near the transition from A class to B class (Fig. S3a–b). This suggests likely underestimation of morbidity expansion in *daf-2(e1370)* in our recent study, in which canonical A and A’ were not distinguished (Zhang and Gems, 2026).

We therefore assessed how including A’-span within our definition of G-span changes the effect of DAF-2 AID from L4 on relative morbidity. As expected, with the A’-adjusted definitions of H-span and G-span, DAF-2 AID caused a smaller increase in H-span and greater increase in G-span, compared to the unadjusted definition (where H-span = canonical A-span + A’-span) (Fig. 4g). Notably, under the A’-adjusted definitions, morbidity is no longer compressed but more proportionally scaled (Fig. 4h). Similarly, A’-adjustment of H-span and G-span definitions increased the contribution of G-span to *daf-2(e1370)* longevity, here amplifying the existing expansion of relative morbidity, compared to N2 (Fig. S3c–d).

In conclusion, this analysis shows using two definitions of gerospan, including the more inclusive A’-adjusted definition, that unlike *daf-2(e1370)* mutation, stronger, adult-restricted, post-translational IIS reduction by DAF-2 AID does not expand relative morbidity. That DAF-2 AID from L4 markedly extends H-span with little expansion of the existing G-span indicates a reduced biological aging rate throughout most of adulthood. Notably, the reduction of only *α* by DAF-2 AID from L4 (Fig. 1e–f) also agrees with our recent finding that reduction of *α* but not *β* is driven by healthspan rather than gerospan expansion, further supporting the conclusion that reduced *α* generally corresponds to reduced biological aging rate (Zhang and Gems, 2026). Taken together, this work shows how DAF-2 AID from the start of adulthood produces a strong, inter-individually homogeneous extension of healthy lifespan, with little increase in morbidity.

### Restoring IIS during advanced senescence extends lifespan and healthspan

Finally, we explored further the effects of IIS in the terminal stages of the aging process. It was previously shown that initiating DAF-2 AID at very advanced ages (e.g. days 20– 25) can extend lifespan (Venz et al., 2021), suggesting possible rejuvenation effects (Molière et al., 2024). However, our longitudinal methodology revealed that even at such advanced ages many individuals are still in the healthiest (A) locomotory class (Fig. 2a–b, h–i), due to slower aging rates in these longer-lived population members. Thus, at the time of treatment they are relatively youthful, and not truly decrepit. Or in other words, many animals on days 20–25 are advanced in chronological but not biological age. It is therefore possible that life extension from very late-life DAF-2 AID reflects anti-aging responses in these biological youthful individuals, rather than anti-aging effects in decrepit individuals.

To test whether DAF-2 AID has any effect on B and C class animals, which are biologically older, we assessed the post-treatment survival of animals in which DAF-2 AID started while they were in B or C class (Fig. 5a). As suspected, no increase in lifespan was seen but, notably, a weak but consistent trend of lifespan *reduction* was detected (Fig. 5a). This raises the possibility that reducing IIS at advanced biological ages is not merely ineffective at extending lifespan, but even detrimental to health.

**Fig. 5.**
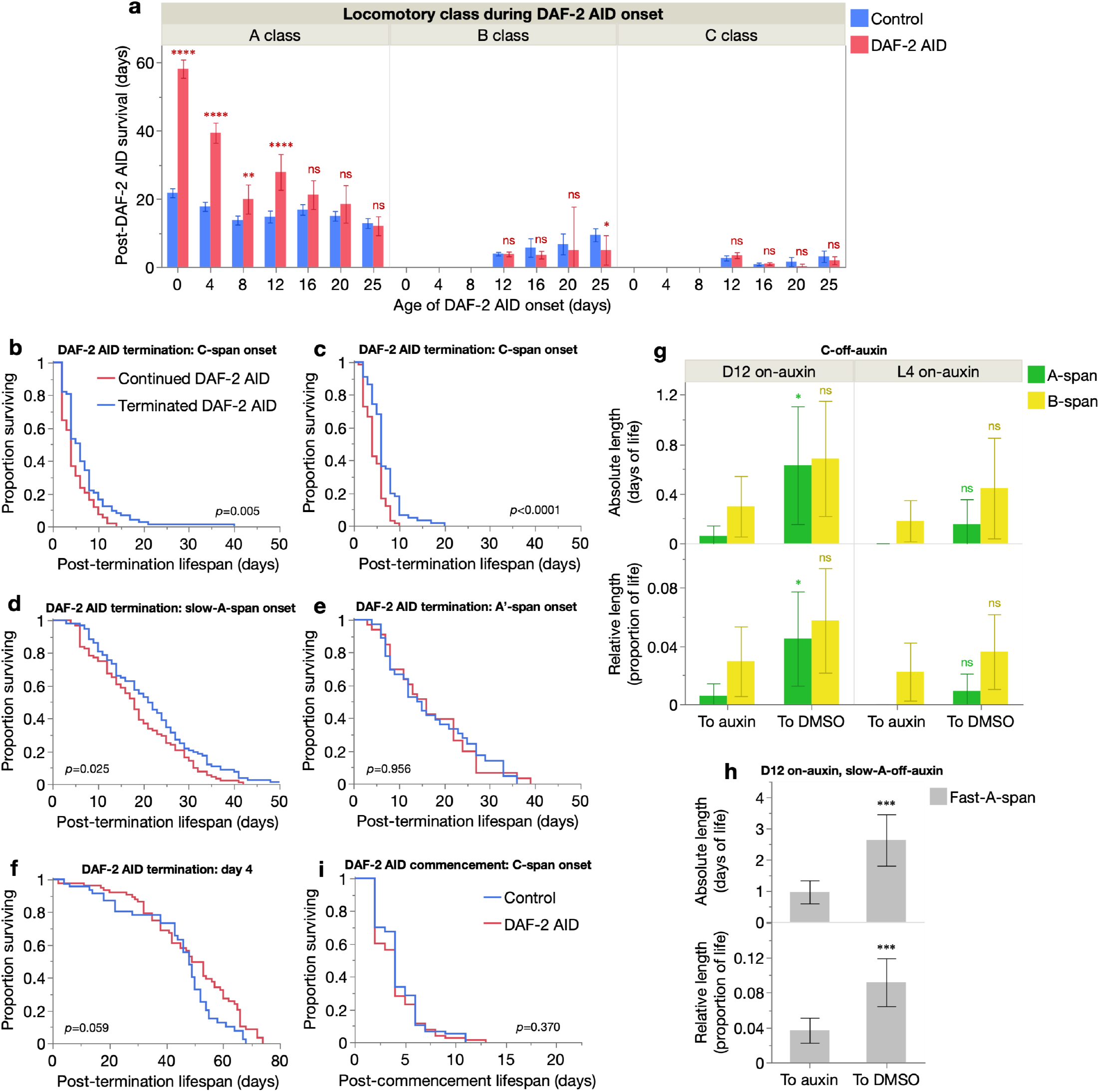
Late-life impacts of IIS on lifespan and locomotory health. (**a**) Effect of auxin treatment on mean days survived after auxin commencement for different locomotory classes (A, B or C), for each age of auxin commencement. 95% confidence intervals shown, and statistical comparisons between DMSO and auxin cohorts made by two-tailed Student’s t-tests; ns *p* > 0.05, * *p* ≤ 0.05, ** *p* ≤ 0.01, *** *p* ≤ 0.001, **** *p* ≤ 0.0001. Data from the pool of Trials 1–5, and censored individuals were excluded (sample sizes in Table S1). (**b**–**f**) Kaplan-Meier survival curves of RAF2181 animals started on auxin from day 12 (**b, d, e**) or L4 (**c, f**) and kept on auxin (red) or moved to DMSO (blue) at the onset of C (**b**– **c**), slow-A (**d**) or A’ (**e**) class locomotion, or on day 4 of adulthood (**f**). Survival curves were compared by log-rank tests, with *p*-values labelled. (**g**–**h**) Effect of moving RAF2181 animals off auxin (to DMSO) at the onset of C (**g**) or slow-A (**h**) class locomotion, on absolute and relative durations of A-span and B-span (**g**) and fast-A-span (**h**). Statistical comparisons between auxin and DMSO cohorts were made by two-tailed Student’s t-tests. (**i**) Kaplan-Meier survival curves of RAF2181 animals started on auxin or DMSO from the onset of C class locomotion. All data are from the pool of two trials, total[censored]: **b, g left** (Continued: *n*=68[1], Terminated: *n*=73[0]); **c, g right** (Continued: *n*=66[0], Terminated: *n*=66[1]); **d, h** (Continued: *n*=92[0], Terminated: *n*=94[2]); **e** (Continued: *n*=78[0], Terminated: *n*=77[0]); **i** (Control: *n*=77[0], DAF-2 AID: *n*=78[0]).

These results suggest that the level of IIS that is optimal for longevity and health increases in the final stages of life. To test this directly, we examined the effect on lifespan of terminating DAF-2 AID in elderly individuals. Here, DAF-2 AID was initiated in A class animals on day 12, and when they entered C-span, DAF-2 AID was either continued until death (control) or terminated (by transfer to DMSO-only plates, until death). Subsequent lifespan was then compared in the two treatments. This revealed an increase in post-treatment lifespan of populations in which DAF-2 AID was terminated (+45% mean post-treatment lifespan, log rank *p*=0.005; Fig. 5b).

Commencing DAF-2 AID from L4 yielded the same result (+48% mean post-treatment lifespan, log rank *p*<0.0001; Fig. 5c). These magnitudes of post-transfer survival increase (from DAF-2 AID commenced from L4 versus day 12) were not significantly different (Cox proportional hazards likelihood-ratio test *p*=0.70), i.e. late-life IIS effects on lifespan were unaffected by the duration of prior knock-down of IIS. Therefore, life-extension by earlier-life IIS reduction and its later-life reversal are additive.

Next, we wondered how early on in the aging process DAF-2 AID cessation can increase lifespan. To investigate this, we assessed the effects of terminating DAF-2 AID at several points during early senescence: onset of A’ locomotion (delayed sinusoidal locomotion), onset of slow-A locomotion (slow sinusoidal locomotion, delayed or not; see Methods), and on day 4 of adulthood (when all individuals exhibit non-delayed and fast sinusoidal locomotion). Terminating DAF-2 AID at slow-A onset also increased mean post-treatment lifespan, but by a smaller magnitude (+19%, log rank *p*=0.025; Fig. 5d), while terminating it at A’ onset did not (+1.5%, log rank *p*=0.956; Fig. 5e). As expected, termination of DAF-2 AID (started from L4) even earlier, on day 4 of adulthood, also did not extend mean post-treatment lifespan (−10%, log rank *p*=0.059; Fig. 5f).

Strikingly, this termination of DAF-2 AID on day 4 only weakly limited the lifespan-extension of DAF-2 AID treatment from L4 (from +154% to +132% of mean lifespan, and +194% to +189% of median lifespan) (Fig. 5f, S4), showing that memory of IIS reduction during the first few days of adulthood is retained throughout life. This is consistent with lasting life-shortening effects of transient increases in IIS during early adulthood in *C. elegans* and fruit flies (Dobson et al., 2017). Taken together, these results indicate that the transition from beneficial to deleterious effects of DAF-2 AID occurs relatively early in aging, in the later stages of A-span.

We then assessed whether restoring IIS in later life also extends healthspan. Notably, regardless of DAF-2 AID initiation age (L4 or day 12), animals removed from treatment at the onset of C class locomotion showed a consistent trend of increased reversal to A and B classes (Fig. 5g), indicating a modest reversal of the aging-related loss of locomotory capacity. Similarly, animals with DAF-2 AID stopped at the onset of slow-A locomotion showed a strong increase in both absolute and relative fast-A-span (fast sinusoidal locomotion, i.e. faster than slow-A; see Methods) post-transfer (Fig. 5h), revealing a robust reversal of locomotory speed decline. These results show that it is possible to rejuvenate certain aging-related traits by late-life IIS modulation, but by restoring/increasing IIS rather than by decreasing it (Molière et al., 2024).

Finally, to further explore the anti-aging effect of late-life IIS, we initiated DAF-2 AID at C-span onset. Notably, this did not extend lifespan, but did not clearly reduce it either (−8.1% mean lifespan, log rank *p*=0.370; Fig. 5i). This weak shortening could potentially reflect reduced efficacy of DAF-2 AID in decrepit animals (i.e. inefficient IIS reduction).

## Discussion

In this study we have applied a longitudinal, individual-focused approach to understand the role of insulin/IGF-1 signaling (IIS) in aging over the life course. This approach enables one to distinguish between and interrelate biological and demographic aging phenomena, given that the latter emerge from the presence of differences in the aging process between population members. We have previously employed this approach to explain the biological basis of the Gompertz mortality model parameters in long-lived *C. elegans* (Zhang and Gems, 2026), and of inverse Strehler-Mildvan correlations between the Gompertz parameters (Zhang et al., 2026).

In the present study we used this approach to capture the wide variability in aging trajectories between isogenic population members, and the resulting variability in their responses to altered IIS. We describe intrinsic inter-individual differences in aging rate within control populations, where longer-lived individuals age more slowly, resulting in the emergence of biological age variation with increasing chronological age. This explains individually-heterogeneous lifespan responses to later-age IIS reduction (Venz et al., 2021), given that the life-extending effects of IIS reduction are diminished with increasing biological age. Indeed, commencing DAF-2 AID from L4, prior to aging-related divergence in individual biological age, yielded homogeneous lifespan increases across population members, shifting the survival curve rightward. This contrasts with effects of IIS reduction by *daf-2* mutation and RNAi, which typically stretch the survival curve, reflecting increased lifespan variation (Johnson, 1990, Dillin et al., 2002, Samuelson et al., 2007, Zhang and Gems, 2026).

Why might these methods of IIS reduction cause such different demographic effects? One possibility is that *daf-2* mutants, which carry partial reduction-of-function mutations (Gems et al., 1998), exhibit inter-individually variable mutant expressivity. Similarly, the efficacy of RNAi varies between individuals, for instance due to differences in dsRNA ingestion. These individually variable reductions of IIS, including during development in *daf-2* mutants, could explain the greater lifespan variation resulting from *daf-2* mutation and RNAi than DAF-2 AID (Fig. 6a). Notably, DAF-2 AID rapidly reduces DAF-2 protein abundance by ∼40% when started from L4, and increases lifespan more than the widely-used, canonical *daf-2(e1370)* mutant (Venz et al., 2021, Zhang et al., 2022). Similarly, *daf-2* RNAi produces weaker lifespan increases, whose magnitude reduces with age faster than that of DAF-2 AID (Dillin et al., 2002). Clearly, the greater magnitude of IIS reduction by DAF-2 AID is closer to the ideal level for life-extension, thus maximising lifespan responses, at least during A-span.

**Fig. 6.**
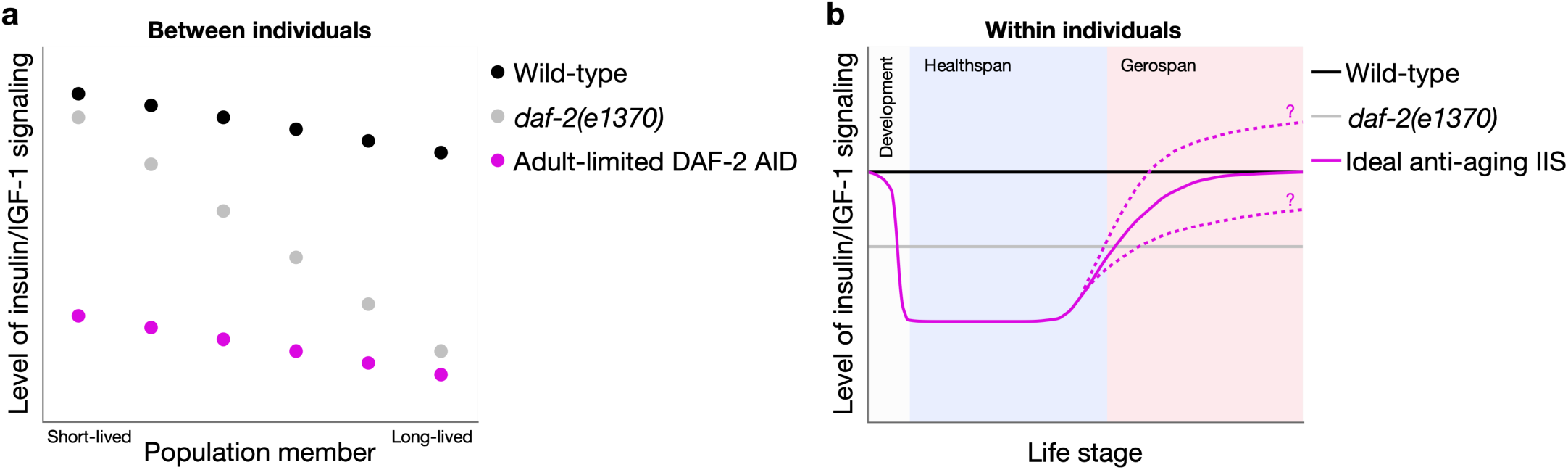
Summary scheme of insulin/IGF-1 signaling effects between and within individuals. (**a**) A model of inherent inter-individual variation in IIS level, amongst isogenic individuals of wild-type, *daf-2(e1370)* and DAF-2 AID-treated nematode populations. *daf-2(rf)* mutations such as *daf-2(e1370)* reduce IIS variably (producing variable increases in lifespan), whereas DAF-2 AID from the start of adulthood reduces IIS more homogeneously and strongly (producing larger and less variable increases in lifespan). (**b**) A model of how the optimal level of IIS for health and longevity changes over the life course, within any population member. During development, a sufficient level of IIS must be maintained to avoid larval arrest, although whether a modest reduction prior to adulthood could have anti-aging effects remains unknown. From the onset of adulthood (in this study treated from L4), a low level of IIS (lower than in *daf-2(e1370)*) promotes health and longevity, but towards the end of healthspan and during gerospan, higher levels of IIS are optimal. Whether the end of life optimal IIS levels eventually exceed wild-type levels, or remain below it, remains unclear (indicated by question marks; see Discussion).

Our longitudinal rather than cross-sectional design revealed a mid-life reversal of DAF-2 AID effects on aging, towards the end of A-span (the duration of sinusoidal locomotory capacity) (Fig. 6b). After this stage, continued IIS reduction becomes life- and health-limiting, and its termination extends lifespan and even modestly rejuvenates locomotory health. We show that previous reports to the contrary, of lifespan and health benefits of DAF-2 AID initiated in late life (Venz et al., 2021, Molière et al., 2024), are attributable to effects on chronologically aged but biologically youthful population members (i.e. with a slower biological aging rate). This highlights the value of characterizing inter-individual variation when studying intra-individual biological traits, particularly those as heterogeneous as the aging process. Our demonstration raises the possibility that other late-age interventions, including in other species, could be affected by similar confounds and should therefore be interpreted with care.

One possible interpretation of these effects of IIS on aging is that they reflect a form of antagonistic pleiotropy (AP), wherein IIS promotes aging in earlier life but protects against it in later life. Evolutionary theory argues that AP traits can evolve where they confer reproductive fitness benefits of greater magnitude than that of fitness detriments in later life (i.e. aging) (Williams, 1957). Here IIS presents a complex picture. Selection for elevated IIS in early life accelerates development, growth and reproductive output, most obviously by preventing dauer larva formation (Riddle et al., 1981), at the cost of accelerated aging in later life (i.e. AP) (Jenkins et al., 2004, Chen et al., 2007, Maklakov et al., 2017). Our findings could imply that at advanced biological ages, wild-type levels of IIS are optimal for viability; that is, they promote aging throughout much of adulthood, but suppress it near the end of life, revealing a new facet of AP in *daf-2*. By extension, an intriguing question is whether these late-life anti-aging effects could be enhanced by increasing IIS above wild-type levels (Fig. 6b). Supporting this possibility is the observation that dietary glucose supplementation, which increases IIS and ordinarily shortens *C. elegans* lifespan (Lee et al., 2009), can extend it when started in mid-life (day 5 of adulthood), although this was true only of chemically-sterilized animals (Beaudoin-Chabot et al., 2022).

However, a possible alternative interpretation is that the degree to which DAF-2 AID reduces IIS is too severe for decrepit elderly nematodes, in which milder IIS reductions are optimal for health and survival (Fig. 6b). In this scenario, wild-type *daf-2* function promotes aging at all stages of adulthood, i.e. the life-extending effect of cessation of DAF-2 AID in late life is not indicative of AP. Either way, our findings demonstrate a later-life increase in the optimal level of IIS for health and longevity.

We also demonstrate that IIS reduction by DAF-2 AID extends lifespan primarily through increases in healthspan, with little increase in the duration of morbidity (gerospan). One possibility is that this lack of gerospan extension reflects, to some degree, the acceleration of aging by severe IIS reduction from late-A-span onwards, which we found in this study. Imagining for a moment an equivalent form of intervention in humans, this could present a somewhat unwholesome prospect: a treatment that extends the healthy portion of life but achieves compression of morbidity partly by accelerating aging in the frail elderly.

Notably, these greater healthspan than gerospan increases caused morbidity compression, something not previously observed in IIS-reducing interventions. Using *daf-2* mutants and different locomotory healthspan definitions, earlier studies typically reported gerospan-driven longevity (i.e. morbidity expansion) (Huang et al., 2004, Bansal et al., 2015, Podshivalova et al., 2017, Churgin et al., 2017, Newell Stamper et al., 2018, Zhang and Gems, 2026) or proportional increases in healthspan and gerospan (Hahm et al., 2015, Statzer et al., 2022). Here we show that stronger, adult-restricted, post-translational IIS reduction can extend lifespan and healthspan together. Our definition of healthspan describes locomotory capacity (maximum response to a stimulus), using a standard measure of locomotory aging (Hosono et al., 1980, Herndon et al., 2002) that integrates the aging-related health status of multiple bodily systems. Indeed, modifying our definition of healthspan by excluding from it A’ locomotion (which we characterized here as an early senescent stage) suggested at most a proportional scaling of healthspan and gerospan, rather than morbidity expansion. Either way, our findings show that IIS reduction can extend lifespan through robust increases in healthspan.

The compression of human morbidity has long been an aspiration of aging research (Fries, 1980, Kannisto, 2000, Seaman et al., 2020). This is often envisioned as resulting from the extension of healthspan against a fixed lifespan (i.e. morbidity compression without life-extension), a view motivated by biological and ethical considerations. However, our findings show that in a relatively simple animal it is possible to increase maximum lifespan without morbidity expansion, suggesting at least the possibility of similar biodemographic changes in humans.

In conclusion, our longitudinal study reveals roles of IIS across the nematode life course as well as between isogenic individuals. During early adulthood, IIS promotes aging, such that its reduction can greatly extend lifespan and healthspan, thereby compressing morbidity. However, in later adulthood, higher levels of IIS are optimal to protect against aging, such that its reduction can shorten lifespan and healthspan. Notably, these intra-individual roles of IIS vary between isogenic individuals, proceeding at different rates according to individual-specific aging rates. Importantly, these findings were attainable through the combined study of these biological traits and their inter-individual variation.

## Materials and Methods

### *C. elegans* culture and strains

*C. elegans* were maintained at 20°C using standard protocols (Brenner, 1974), on Nematode Growth Medium (NGM, prepared using BactoPeptone) plates seeded with an *Escherichia coli* OP50 bacterial food source 2 days prior to use, and without the addition of antibiotics, or 5-fluoro-2-deoxyuridine (FUDR) which is sometimes used to block progeny production. Nematode strains used were: N2 (wild-type, hermaphrodite stock (Zhao et al., 2019)), GA1960 *daf-2(e1368) III*, GA1928 *daf-2(e1370) III*, and RAF2181 *ieSi57 II; daf-2(bch40) III* (Venz et al., 2021).

### Age-specific DAF-2 AID

All age-specific DAF-2 AID (auxin-inducible degradation) cohorts (comprised of RAF2181 animals) were cultured on DMSO plates from L4 until their age of auxin commencement (L4, or days 4, 8, 12, 16, 20 or 25 of adulthood), when they were then transferred to auxin plates. DMSO plates were prepared one day before use, by topically adding 5 μL of 100% DMSO to the side of the *E. coli* lawn of each 2 mL NGM-filled well of 24-well tissue culture plates. Similarly, 24-well auxin plates were prepared by topically adding 5 μL of 400 mM auxin (indole-3-acetic acid, Sigma #I3750; dissolved in DMSO and stored long-term at −25°C, or 4°C for up to 1 week) to each well, one day prior to use. Auxin crystals initially precipitate on the media surface, but redissolve by the next day. One worm was placed in each well to facilitate their longitudinal culture and study, and animals were transferred to fresh 24-well plates before media desiccation. Lifespan and locomotory capacity class (see below) were scored every 2– 3 days. Individuals were censored if they died from desiccation on the plate wall, from rupture of internal organs through the vulva, from internal hatching of larvae, and if they were unable to be located.

### Quantification of locomotory decline during aging

Locomotory health class was scored by classifying individuals into one of three main classes, adapted from earlier systems (Hosono et al., 1980, Herndon et al., 2002): A – sinusoidal locomotion; B – non-sinusoidal locomotion; C – no locomotion. To accurately determine locomotory class, animals were first gently touched (repeatedly, where necessary), on the tail with a platinum wire worm pick for up to 20 seconds to induce an escape response that reveals movement capacity (rather than food-related preference (Hahm et al., 2015)), and additionally on the head as a final check. Animals had to travel at least one body length’s distance for sinusoidal (S-shaped) locomotion to be registered; any irregularities to S-shaped locomotion or failure to travel at least one body length’s distance within 20 seconds resulted in classification as B class. C class individuals were scored if animals could not travel at all and exhibited only stationary movements, usually of the head and/or tail. Very old C class animals were distinguished from dead animals by application of a greater wire force, which can reveal residual movement capacity.

Sub-classes of A class animals were distinguished in some experiments (Fig. 4, 5d–e, h, S3): (1) A’ animals, which begin sinusoidal locomotion only after 5 seconds of a single touch of the tail with the platinum wire, compared to (2) canonical A animals which begin sinusoidal locomotion within 5 seconds of the touch, (3) slow-A animals, which travel noticeably slower by eye than more youthful (4) fast-A animals. A class can therefore be split into A’ and canonical A based on locomotory responsiveness, or slow-A and fast-A based on locomotory speed, and individuals can display any combination of responsiveness and speed.

### Late-life termination of DAF-2 AID

Prior to auxin treatment commencement, animals were cultured on standard NGM plates (without DMSO). These experiments were performed in 24-well tissue culture plates, except for day 4 termination of DAF-2 AID (Fig. 5f, S4), which was performed on 60 mm diameter plates containing 10 mL NGM (and 25 μL of 100% DMSO or 25 μL of 400 mM auxin, added topically around the *E. coli* lawn one day prior to use).

### Statistics and software

All statistical tests were performed in JMP Pro (SAS Institute, Inc.), except for Gompertz parameter estimation by maximum likelihood estimation, and assessment of statistical differences between the Gompertz parameters by likelihood ratio tests, which were performed in WinModest (Pletcher, 1999). Censored individuals were included as right censors in all Kaplan-Meier and WinModest analyses, and excluded from other analyses. Specific statistical tests and associated methodological details are described in the respective figure/table captions. Notation of statistical significance in all figures is as follows: *p* > 0.05, * *p* ≤ 0.05, ** *p* ≤ 0.01, *** *p* ≤ 0.001, **** *p* ≤ 0.0001.

## Data availability

Raw data is provided as a Source Data file.

## Supplementary Material

Supplementary Figures 1–4, Supplementary Tables 1–3.

## Funding information

This work was supported by a Wellcome Trust Investigator Award (215574/Z/19/Z) to DG.

## Acknowledgments

We thank Yiran Zhang for minor research contributions and Adrian Moliere for comments on the manuscript. Some strains were provided by the Caenorhabditis Genetics Center, which is funded by the NIH Office of Research Infrastructure Programs (P40 OD010440).

## Author contributions

DG supervised the project. DG and BZ conceived the project, designed the experiments and data analysis, and wrote the manuscript, with assistance from CYE. BZ, KCH, RB, MC, XW, HC, MK, AZ and CN performed the experiments and BZ analyzed the data.

## Conflict of interest

With no relation to the present manuscript, CYE is a co-founder and shareholder of Avea Life AG and Lichi3 GmbH and is employed by Novartis. All other authors declare no conflicts of interest.

