## Supplementary Data for "Strong inhibition of insulin/IGF-1 signaling in early-mid adulthood compresses morbidity, but in later life accelerates aging"

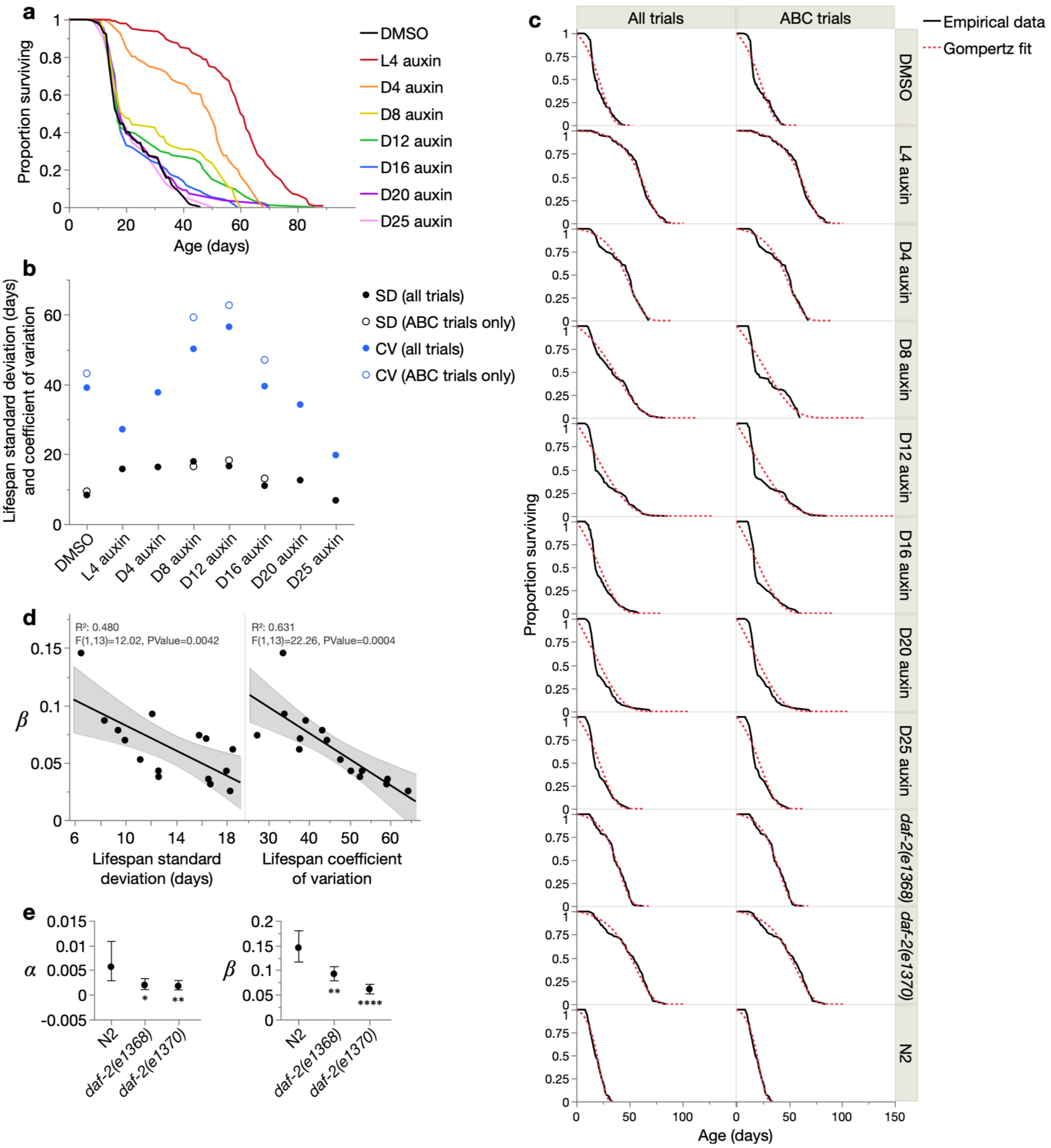


**Fig. S1.** (**a**) Kaplan-Meier survival curves for the age-specific DAF-2 AID and DMSO-only control cohorts (pool of trials in which locomotory capacity was scored; see Table S1). (**b**) Standard deviation (SD) and coefficient of variation of lifespan (CV) of the control and DAF-2 AID cohorts (including only individuals dying *after* the day DAF-2 AID commencement), for the pool of all trials and only trials in which locomotory capacity was scored. Censored individuals were excluded. (**c**) Overlay of empirical survival curves and Gompertz survival fits for each cohort, including two *daf-2* mutants and N2, for the pool of all trials and only trials in which locomotory capacity was scored. (**d**) Least-squares linear regressions between the Gompertz parameter $\beta$ and lifespan standard deviation (left) and lifespan coefficient of variation (right), displaying 95% confidence regions and F-test statistics. (**e**) Effect of *daf-2(e1368)* and *daf-2(e1370)* mutation on the Gompertz parameters $\alpha$ and $\beta$ (relative to the N2 wildtype). Statistical significance of parameter differences were assessed by likelihood ratio tests, displaying 95% confidence intervals; ns $p > 0.05$, * $p \leq 0.05$, ** $p \leq 0.01$, *** $p \leq 0.001$, **** $p \leq 0.0001$.

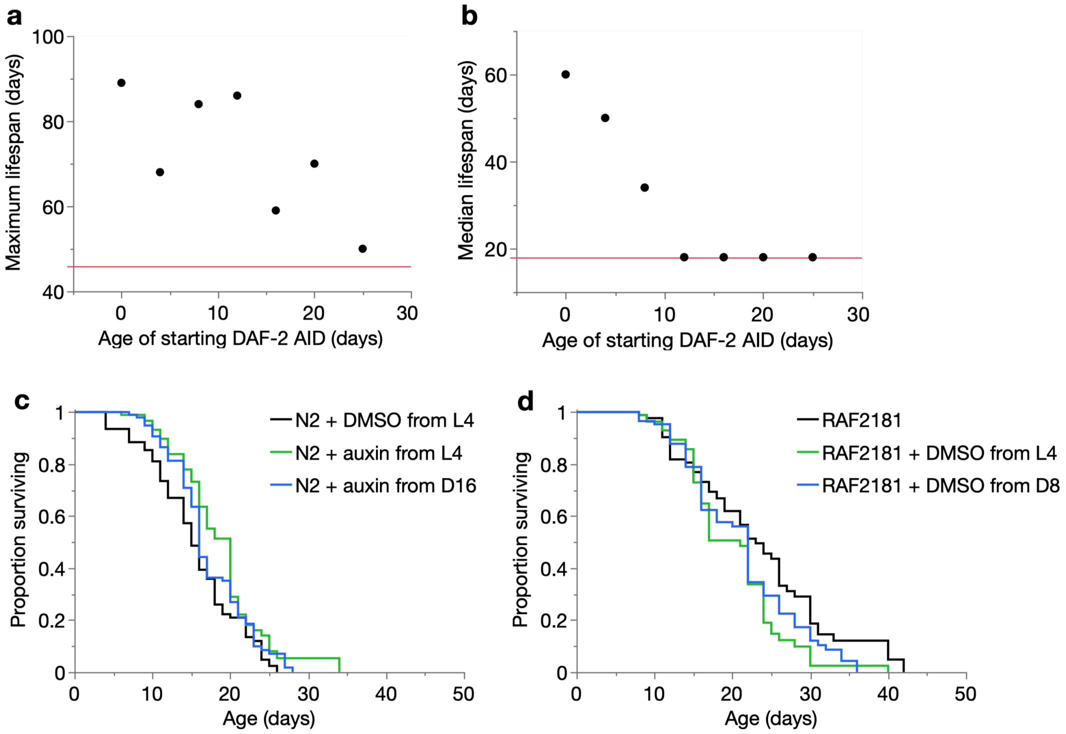


**Fig. S2.** (**a**–**b**) Effect of age of commencement of auxin administration on (**a**) maximum and (**b**) median lifespan (from the pool of all trials). The horizontal red lines (y-axis=46 in **a** and 18 in **b**) denotes the maximum and median lifespans of the DMSO control, respectively. (**c**–**d**) Kaplan-Meier survival curves of (**c**) N2 animals treated with DMSO from L4 onwards (mean lifespan: 15.4 days; *n*=108[25]), or auxin from either L4 (mean lifespan: 18.7 days; *n*=96[24]) or day 16 (mean lifespan: 17.2 days; *n*=108[21]) onwards, and (**d**) RAF2181 animals without DMSO (mean lifespan: 23.5 days; *n*=96[37]), or treated with DMSO from either L4 (mean lifespan: 20.1 days; *n*=96[46]) or day 8 (mean lifespan: 21.3; *n*=96[31]) onwards. Pool of two trials. Sample size notation: total[censored]

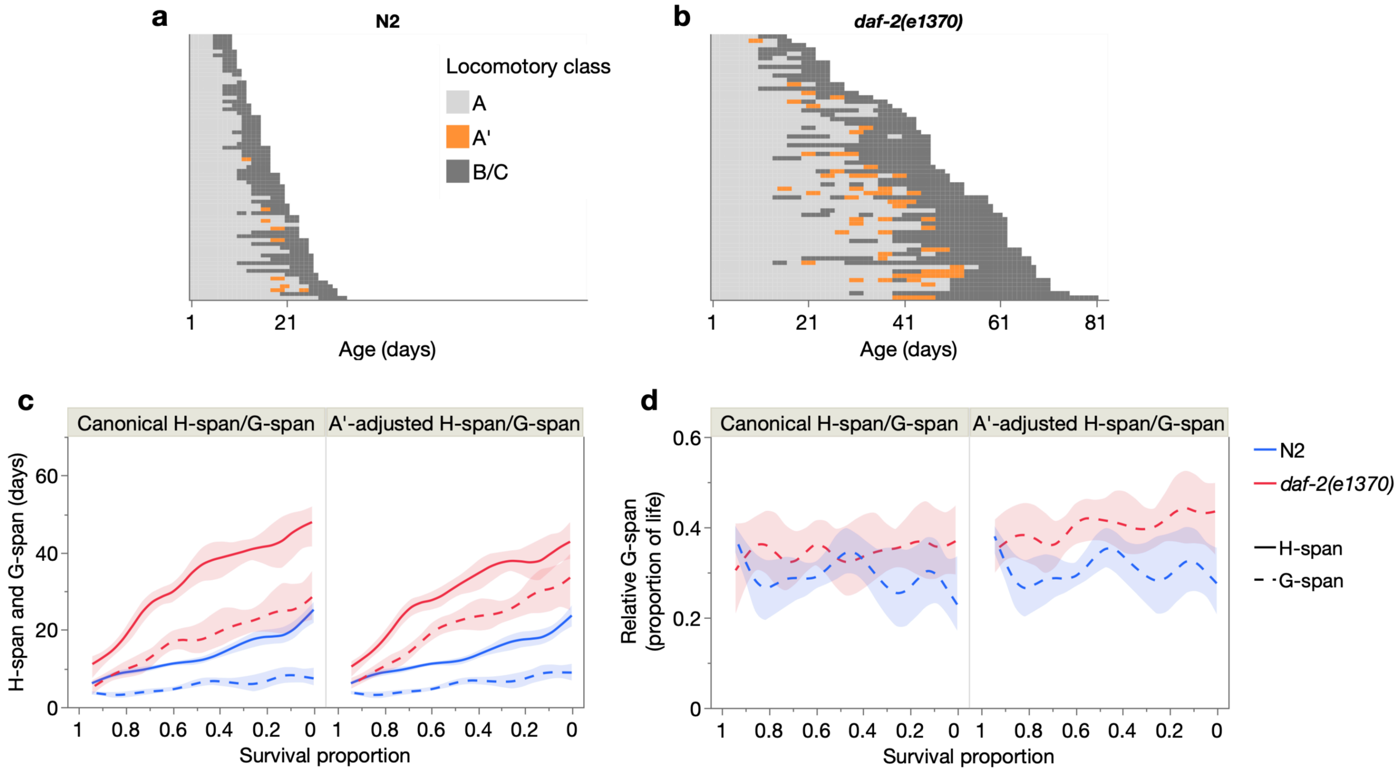


**Fig. S3.** (**a**–**b**) Event history charts of the locomotory class of each individual throughout life, of N2 (**a**) and *daf-2(e1370)* animals (**b**). Each row represents one individual. (**c**–**d**) Effect of *daf-2(e1370)* on (**c**) absolute H-span and G-span and (**d**) relative G-span, using either the canonical definitions (H-span = A-span, G-span = B+C-span) or A'-adjusted definitions (H-span = canonical A-span, G-span = A'+B+C-span). These relationships were fit with a smoother (spline method, lambda=0.05) displaying 95% confidence regions. All panels (**a**–**d**) are from the pool of Trials 3 and 5, in which A' was scored (sample sizes in Table S1).

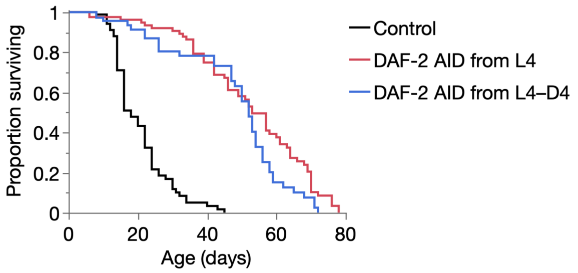


**Fig. S4.** Kaplan-Meier survival curves of nematodes treated with only DMSO from L4 onwards (Control; *n*=96[32]), auxin from L4 onwards (red cohort; *n*=123[59]), or auxin from L4 until day 4 of adulthood and then followed by DMSO until death (blue cohort; *n*=102[60]). Sample size notation: total[censored]; note that DAF-2 AID from L4 produces a high rate of censors due to bagging. Pool of two trials.
