## Supplementary Tables for "Strong inhibition of insulin/IGF-1 signaling in early-mid adulthood compresses morbidity, but in later life accelerates aging"

| | | | | | Lifespan | | $\alpha$ | | $\beta$ | |
|---|---|---|---|---|---|---|---|---|---|---|
| Cohort | No. dead/ censors | Mean lifespan (days since L4) | $\alpha$ | $\beta$ | % change vs. DMSO | $p$ vs. DMSO (Log-Rank) | % change vs. DMSO | $p$ vs. DMSO (LRT) | % change vs. DMSO | $p$ vs. DMSO (LRT) |
| RAF2181 + DMSO | **[1–7] 282/12** | **21.7** | 0.0107 | 0.0869 | | | | | | |
| | **[1–5] 197/12** | **22.4** | 0.0113 | 0.0784 | | | | | | |
| | [1] 65/0 | 23.6 | | | | | | | | |
| | [2] 36/0 | 22.3 | | | | | | | | |
| | [3] 36/0 | 21.4 | | | | | | | | |
| | [4] 25/11 | 23.9 | | | | | | | | |
| | [5] 35/1 | 19.6 | | | | | | | | |
| | [6] 20/0 | 21.3 | | | | | | | | |
| | [7] 65/0 | 19.9 | | | | | | | | |
| RAF2181 + L4 auxin | **[1–7] 137/45** | **58.6** | 0.0006 | 0.0740 | **169.9** | **<0.0001** | -94.7 | 1.67E-23 | -14.9 | 0.0848 |
| | **[1–5] 137/45** | **58.6** | 0.0006 | 0.0740 | **161.2** | **<0.0001** | -95.0 | 1.36E-21 | -5.6 | 0.5883 |
| | [1] 47/0 | 60.0 | | | 153.8 | <0.0001 | | | | |
| | [2] 24/12 | 63.6 | | | 185.4 | <0.0001 | | | | |
| | [3] 26/10 | 58.8 | | | 175.1 | <0.0001 | | | | |
| | [4] | | | | | | | | | |
| | [5] 40/23 | 54.0 | | | 175.0 | <0.0001 | | | | |
| | [6] | | | | | | | | | |
| | [7] | | | | | | | | | |
| RAF2181 + D4 auxin | **[1–7] 120/16** | **44.6** | 0.0018 | 0.0711 | **105.3** | **<0.0001** | -82.9 | 5.33E-10 | -18.1 | 0.0572 |
| | **[1–5] 120/16** | **44.6** | 0.0018 | 0.0711 | **98.7** | **<0.0001** | -83.8 | 1.48E-09 | -9.2 | 0.4144 |
| | [1] 64/0 | 43.1 | | | 82.5 | <0.0001 | | | | |
| | [2] 32/4 | 47.6 | | | 113.6 | <0.0001 | | | | |
| | [3] | | | | | | | | | |
| | [4] 24/12 | 43.8 | | | 83.5 | <0.0001 | | | | |
| | [5] | | | | | | | | | |
| | [6] | | | | | | | | | |
| | [7] | | | | | | | | | |
| RAF2181 + D8 auxin | **[1–7] 161/12** | **36.5** | 0.0078 | 0.0429 | **68.0** | **<0.0001** | -27.5 | 0.1309 | -50.7 | 1.71E-10 |
| | **[1–5] 60/12** | **30.3** | 0.0133 | 0.0358 | **35.2** | **<0.0001** | 17.4 | 0.5883 | -54.3 | 2.04E-05 |
| | [1] | | | | | | | | | |
| | [2] 34/2 | 29.8 | | | 33.9 | 0.007 | | | | |
| | [3] | | | | | | | | | |
| | [4] 26/10 | 31.0 | | | 29.8 | 0.2345 | | | | |
| | [5] | | | | | | | | | |
| | [6] 43/0 | 44.9 | | | 111.4 | <0.0001 | | | | |
| | [7] 58/0 | 37.1 | | | 86.8 | <0.0001 | | | | |
| RAF2181 + D12 auxin | **[1–7] 183/2** | **28.3** | 0.0169 | 0.0314 | **30.4** | **<0.0001** | 57.5 | 0.0151 | -63.9 | 1.11E-16 |
| | **[1–5] 94/2** | **28.4** | 0.0190 | 0.0254 | **26.7** | **<0.0001** | 68.1 | 0.0306 | -67.6 | 6.94E-11 |
| | [1] | | | | | | | | | |
| | [2] 34/2 | 29.3 | | | 31.6 | 0.0179 | | | | |
| | [3] | | | | | | | | | |
| | [4] | | | | | | | | | |
| | [5] 60/0 | 27.9 | | | 42.2 | 0.0008 | | | | |
| | [6] 20/0 | 38.7 | | | 82.1 | <0.0001 | | | | |
| | [7] 69/0 | 25.2 | | | 26.6 | 0.0035 | | | | |
| RAF2181 + D16 auxin | **[1–7] 178/2** | **23.4** | 0.0160 | 0.0528 | **7.9** | **0.0379** | 49.5 | 0.0362 | -39.3 | 3.87E-06 |
| | **[1–5] 94/2** | **23.8** | 0.0183 | 0.0430 | **6.0** | **0.0665** | 61.8 | 0.0500 | -45.1 | 0.0001 |
| | [1] | | | | | | | | | |
| | [2] 46/2 | 22.9 | | | 2.9 | 0.6111 | | | | |
| | [3] | | | | | | | | | |
| | [4] | | | | | | | | | |
| | [5] 48/0 | 24.6 | | | 25.5 | 0.0029 | | | | |
| | [6] 23/0 | 22.5 | | | 6.0 | 0.2385 | | | | |
| | [7] 61/0 | 23.2 | | | 16.8 | 0.0049 | | | | |
| RAF2181 + D20 auxin | **[1–7] 67/5** | **24.8** | 0.0191 | 0.0378 | **14.0** | **0.055** | 77.8 | 0.0207 | -56.5 | 6.05E-09 |
| | **[1–5] 67/5** | **24.8** | 0.0191 | 0.0378 | **10.4** | **0.2332** | 68.6 | 0.0455 | -51.8 | 8.38E-06 |
| | [1] | | | | | | | | | |
| | [2] 67/5 | 24.8 | | | 11.2 | 0.3187 | | | | |
| | [3] | | | | | | | | | |
| | [4] | | | | | | | | | |
| | [5] | | | | | | | | | |
| | [6] | | | | | | | | | |
| | [7] | | | | | | | | | |
| RAF2181 + D25 auxin | **[1–7] 90/5** | **22.5** | 0.0129 | 0.0697 | **3.5** | **0.4172** | 20.1 | 0.4693 | -19.8 | 0.0842 |
| | **[1–5] 90/5** | **22.5** | 0.0129 | 0.0697 | **0.1** | **0.9712** | 13.9 | 0.6258 | -11.1 | 0.4084 |
| | [1] | | | | | | | | | |
| | [2] 90/5 | 22.5 | | | 0.8 | 0.9122 | | | | |
| | [3] | | | | | | | | | |
| | [4] | | | | | | | | | |
| | [5] | | | | | | | | | |
| | [6] | | | | | | | | | |
| | [7] | | | | | | | | | |
| *daf-2(e1368)* | **[1–7] 127/30** | **36.4** | 0.0020 | 0.0926 | **67.6** | **<0.0001** | -81.6 | 9.50E-10 | 6.5 | 0.5261 |
| | **[1–5] 127/30** | **36.4** | 0.0020 | 0.0926 | **62.3** | **<0.0001** | -82.6 | 2.66E-09 | 18.1 | 0.1349 |
| | [1] 43/0 | 32.7 | | | 38.3 | <0.0001 | | | | |
| | [2] 24/12 | 33.8 | | | 51.9 | 0.0005 | | | | |
| | [3] 26/8 | 37.5 | | | 75.5 | <0.0001 | | | | |
| | [4] | | | | | | | | | |
| | [5] 34/10 | 41.5 | | | 111.7 | <0.0001 | | | | |
| | [6] | | | | | | | | | |
| | [7] | | | | | | | | | |
| *daf-2(e1370)* | **[1–7] 148/12** | **49.6** | 0.0018 | 0.0617 | **128.5** | **<0.0001** | -83.2 | 8.37E-12 | -29.0 | 0.0006 |
| | **[1–5] 148/12** | **49.6** | 0.0018 | 0.0617 | **121.2** | **<0.0001** | -84.1 | 4.78E-11 | -21.3 | 0.0364 |
| | [1] 54/0 | 49.7 | | | 110.2 | <0.0001 | | | | |
| | [2] 33/1 | 48.8 | | | 119.0 | <0.0001 | | | | |
| | [3] 30/6 | 58.8 | | | 174.9 | <0.0001 | | | | |
| | [4] | | | | | | | | | |
| | [5] 31/5 | 40.6 | | | 106.6 | <0.0001 | | | | |
| | [6] | | | | | | | | | |
| | [7] | | | | | | | | | |
| N2 | **[1–7] 69/4** | **19.5** | 0.0057 | 0.1457 | **-10.0** | **0.0137** | -47.1 | 0.0608 | 67.6 | 0.0003 |
| | **[1–5] 69/4** | **19.5** | 0.0057 | 0.1457 | **-12.9** | **0.0021** | -49.8 | 0.0488 | 85.9 | 0.0001 |
| | [1] | | | | | | | | | |
| | [2] | | | | | | | | | |
| | [3] 33/3 | 20.6 | | | -3.9 | 0.4619 | | | | |
| | [4] | | | | | | | | | |
| | [5] 36/1 | 18.5 | | | -5.5 | 0.3457 | | | | |
| | [6] | | | | | | | | | |
| | [7] | | | | | | | | | |

**Table S1. Lifespan and Gompertz parameters of all cohorts.** N2: wild-type, + D*n* auxin: auxin given from day *n* of adulthood (*n* days since L4) until death, [1–7]: pool of all trials, [1–5]: pool of trials in which locomotory function (ABC system) was scored, [*n*]: trial number. Trials 1–3 and 5 were performed by BZ, Trial 4 by BZ and RB, Trial 6 by KH and XW, and Trial 7 by KH. LRT: likelihood ratio test, used to assess statistical significance of differences between Gompertz parameters. Gompertz parameters and associated LRTs were performed for pooled data rather than individual trials, given larger sample sizes of the former.

| | DMSO-treated (control) | | Auxin-treated | | | |
|---|---|---|---|---|---|---|
| Treatment day | No. dead/ censors | Mean lifespan (days since L4) | No. dead/ censors | Mean lifespan (days since L4) | % change vs. DMSO | *p* vs. DMSO; Log-Rank [Wilcoxon] |
| 0 (L4) | **[1–7] 282/12** | **21.7** | **[1–7] 137/45** | **58.6** | **169.9** | **<0.0001** |
| | **[1–5] 197/12** | **22.4** | **[1–5] 137/45** | **58.6** | **161.2** | **<0.0001** |
| | [1] 65/0 | 23.6 | [1] 47/0 | 60.0 | 153.8 | <0.0001 |
| | [2] 36/0 | 22.3 | [2] 24/12 | 63.6 | 185.4 | <0.0001 |
| | [3] 36/0 | 21.4 | [3] 26/10 | 58.8 | 175.1 | <0.0001 |
| | [4] 25/11 | 23.9 | [4] | | | |
| | [5] 35/1 | 19.6 | [5] 40/23 | 54.0 | 175.0 | <0.0001 |
| | [6] 20/0 | 21.3 | [6] | | | |
| | [7] 65/0 | 19.9 | [7] | | | |
| 4 | **[1–7] 282/12** | **21.7** | **[1–7] 120/16** | **44.6** | **105.3** | **<0.0001** |
| | **[1–5] 197/12** | **22.4** | **[1–5] 120/16** | **44.6** | **98.7** | **<0.0001** |
| | [1] 65/0 | 23.6 | [1] 64/0 | 43.1 | 82.5 | <0.0001 |
| | [2] 36/0 | 22.3 | [2] 32/4 | 47.6 | 113.6 | <0.0001 |
| | [3] 36/0 | 21.4 | [3] | | | |
| | [4] 25/11 | 23.9 | [4] 24/12 | 43.8 | 83.5 | <0.0001 |
| | [5] | | [5] | | | |
| | [6] 20/0 | 21.3 | [6] | | | |
| | [7] 65/0 | 19.9 | [7] | | | |
| 8 | **[1–7] 281/12** | **21.8** | **[1–7] 161/12** | **36.5** | **67.7** | **<0.0001** |
| | **[1–5] 196/12** | **22.5** | **[1–5] 60/12** | **30.3** | **34.8** | **<0.0001** |
| | [1] 64/0 | 23.9 | [1] | | | |
| | [2] 36/0 | 22.3 | [2] 34/2 | 29.8 | 33.9 | 0.0070 |
| | [3] 36/0 | 21.4 | [3] | | | |
| | [4] 25/11 | 23.9 | [4] 26/10 | 31.0 | 29.8 | 0.2345 |
| | [5] | | [5] | | | |
| | [6] 20/0 | 21.3 | [6] 43/0 | 44.9 | 111.4 | <0.0001 |
| | [7] 65/0 | 19.9 | [7] 58/0 | 37.1 | 86.8 | <0.0001 |
| 12 | **[1–7] 263/12** | **22.5** | **[1–7] 172/0** | **29.4** | **30.9** | **<0.0001** |
| | **[1–5] 183/12** | **23.2** | **[1–5] 90/0** | **29.2** | **25.5** | **<0.0001** |
| | [1] 61/0 | 24.5 | [1] | | | |
| | [2] 32/0 | 23.7 | [2] 33/0 | 29.8 | 25.9 | 0.0421 |
| | [3] 34/0 | 21.9 | [3] | | | |
| | [4] 24/11 | 24.2 | [4] | | | |
| | [5] 32/1 | 20.4 | [5] 57/0 | 28.8 | 41.1 | 0.0009 |
| | [6] 20/0 | 21.3 | [6] 20/0 | 38.7 | 82.1 | <0.0001 |
| | [7] 60/0 | 20.7 | [7] 62/0 | 26.8 | 29.5 | 0.0010 |
| 16 | **[1–7] 162/7** | **26.7** | **[1–7] 121/0** | **27.8** | **4.3** | **0.1676** |
| | **[1–5] 98/7** | **29.6** | **[1–5] 66/0** | **27.8** | **-5.9** | **0.9082 [0.0032]** |
| | [1] 41/0 | 28.9 | [1] | | | |
| | [2] 15/0 | 33.2 | [2] 24/0 | 30.6 | -7.9 | 0.8429 [0.2290] |
| | [3] 17/0 | 28.7 | [3] | | | |
| | [4] 13/6 | 28.6 | [4] | | | |
| | [5] 12/1 | 29.0 | [5] 42/0 | 26.2 | -9.7 | 0.7318 [0.0623] |
| | [6] 17/0 | 22.2 | [6] 14/0 | 27.3 | 23.0 | 0.0043 |
| | [7] 47/0 | 22.1 | [7] 41/0 | 28.0 | 27.0 | <0.0001 |
| 20 | **[1–7] 104/6** | **30.8** | **[1–7] 25/2** | **37.6** | **22.3** | **0.0029** |
| | **[1–5] 73/6** | **33.0** | **[1–5] 25/2** | **37.6** | **14.2** | **0.0504** |
| | [1] 33/0 | 31.4 | [1] | | | |
| | [2] 13/0 | 35.4 | [2] 25/2 | 37.6 | 6.3 | 0.7750 |
| | [3] 10/0 | 36.0 | [3] | | | |
| | [4] 10/5 | 30.5 | [4] | | | |
| | [5] | | [5] | | | |
| | [6] 10/0 | 23.9 | [6] | | | |
| | [7] 21/0 | 26.0 | [7] | | | |
| 25 | **[1–7] 76/4** | **33.4** | **[1–7] 31/0** | **34.5** | **3.0** | **0.2424** |
| | **[1–5] 62/4** | **34.6** | **[1–5] 31/0** | **34.5** | **-0.5** | **0.6337** |
| | [1] 28/0 | 32.9 | [1] | | | |
| | [2] 11/0 | 37.8 | [2] 31/0 | 34.5 | -8.9 | 0.4062 [0.0815] |
| | [3] 10/0 | 36.0 | [3] | | | |
| | [4] 6/3 | 33.4 | [4] | | | |
| | [5] | | [5] | | | |
| | [6] 3/0 | 27.0 | [6] | | | |
| | [7] 11/0 | 28.4 | [7] | | | |

**Table S2. Lifespan statistics for post-auxin treatment deaths only.** [1–7]: pool of all trials, [1–5]: pool of trials in which locomotory function (ABC system) was scored, [*n*]: trial number. Individuals that died or became censored on or before treatment day (with auxin or DMSO) were excluded from these analyses, such that sample sizes decrease down the table (as treatment day gets later). This affects treatments from day 8 (statistics for the earlier treatments are the same as those in Table S1). Where survival curves intersect each other and exhibit greater changes in early than late mortality, the Wilcoxon test is more appropriate to detect lifespan differences than the Log-Rank test; in such cases, both tests were run (see last column).

| Cohort | Anderson-Darling test statistic | *p* value |
|---|---|---|
| DMSO | 1.6944 | 0.1530 |
| DMSO (ABC trials only) | 1.6564 | 0.1596 |
| L4 auxin | 0.3296 | 0.9230 |
| D4 auxin | 0.8887 | 0.4583 |
| D8 auxin | 0.5927 | 0.6769 |
| D8 auxin (ABC trials only) | 1.5825 | 0.1962 |
| D12 auxin | 1.5331 | 0.1900 |
| D12 auxin (ABC trials only) | 1.5863 | 0.1970 |
| D16 auxin | 1.8288 | 0.1312 |
| D16 auxin (ABC trials only) | 1.8472 | 0.1386 |
| D20 auxin | 1.5850 | 0.1980 |
| D25 auxin | 0.8100 | 0.5175 |
| *daf-2(e1368)* | 0.4498 | 0.8086 |
| *daf-2(e1370)* | 0.7920 | 0.5303 |
| N2 | 0.4499 | 0.8118 |

**Table S3. Anderson-Darling goodness-of-fit test statistics for all cohorts.** Test statistics and *p* values for parametric bootstrap Anderson-Darling tests, to assess goodness-of-fit of all cohorts to the MLE Gompertz distributions. This test assesses overall, tail-weighted fit (as the sum of squared, tail-weighted deviations between these functions across all ages). For each cohort, observed A-D statistics were compared against null distributions generated from 5000 parametric bootstrap populations simulated in JMP under the fitted Gompertz distribution of interest (MLE parameters obtained in WinModest), by generating random survival proportions (Uniform(0,1)) and population sizes matching the observed populations (including censored individuals). Monte Carlo *p* values were computed as the proportion of bootstrap A-D statistics greater than or equal to the observed statistic, applying the Phipson–Smyth correction (addition of 1 to numerator and denominator).
